# Peripheral blood cell immunophenotyping reveals distinct subgroups of inflamed depression

**DOI:** 10.1101/706309

**Authors:** Mary-Ellen Lynall, Lorinda Turner, Junaid Bhatti, Jonathan Cavanagh, Peter de Boer, Valeria Mondelli, Declan Jones, Wayne C. Drevets, Philip Cowen, Neil A. Harrison, Carmine M. Pariante, Linda Pointon, NIMA Consortium, Menna R. Clatworthy, Ed Bullmore

## Abstract

Depression has been associated with increased inflammatory proteins but changes in circulating immune cells are less well defined. We used multi-parametric flow cytometry to investigate 14 subsets of peripheral blood cells in 206 cases of major depressive disorder (MDD) and 77 age- and sex-matched controls. There were significant case-control differences, by univariate and multivariate analysis: cases showed increased immune cell counts, especially neutrophils, CD4^+^ T cells and monocytes, and increased inflammatory proteins (C-reactive protein and interleukin-6). Within-group analysis demonstrated significant association between the severity of depressive symptoms and increased myeloid and CD4^+^ cell counts. MDD cases could be partitioned into two groups by forced binary clustering of cell counts: the inflamed depression group (N=81 out of 206; 39%) had increased monocyte, CD4^+^ and neutrophil counts, increased C-reactive protein (CRP) and interleukin 6 (IL-6), and was more depressed than the uninflamed majority of cases. Relaxing the presumption of a binary classification, data-driven clustering identified four subgroups of MDD cases: two of these subgroups (N=38 and N=100; 67% collectively) were associated with increased inflammatory proteins and more severe depression, but differed from each other in the relative weighting of myeloid and lymphoid cell counts. Case-control and within-group results were robust to statistical control for the potentially confounding effects of age, sex, BMI, recent infection status, and tobacco use. Peripheral blood immunophenotyping can be used to identify a candidate cellular biomarker of inflamed depression, and to further decompose that binary partition, suggesting that there is more than one mechanistic pathway underlying inflamed depression.

**One Sentence Summary:** Two subgroups of depressed cases (about two-thirds of all 206 cases) were identified by peripheral blood biomarker evidence of distinctive cellular immunophenotypes, biased towards the myeloid or lymphoid lineages in different subgroups, but consistently associated with increased blood concentrations of inflammatory proteins and greater severity of depressive symptoms.

## Introduction

Major depressive disorder (MDD) affects 10-15% of the population (*1*) and has substantial negative personal, economic and societal effects. At least one third of patients with depression do not respond to conventional antidepressants (*2*) highlighting the need for new therapeutic approaches. Some patients with MDD have evidence of immune activation. The immune system is comprised of an innate and adaptive arm. The former generates a rapid response to relevant stimuli via soluble components, such as C-reactive protein (CRP) and pro-inflammatory cytokines, and a cellular component, involving myeloid cells such as neutrophils, monocytes and macrophages. The adaptive immune system provides a delayed but more specific response via T and B lymphocytes expressing specific antigen receptors. Elevated levels of circulating inflammatory cytokines (interleukin (IL)-6, IL-1β and tumour necrosis factor-alpha (TNFα)) and CRP have been observed at both transcript and protein level in patients with MDD (*3–7*). Global transcriptomic analyses of whole blood have confirmed an increase in innate immune and infection response pathways coupled with a decrease in transcripts associated with the adaptive immune response in MDD patients compared with controls (*8*). These data raise the possibility that immune dysfunction may contribute to MDD pathogenesis. This thesis is supported by animal studies showing that exposure to pro-inflammatory cytokines results in behaviours that have similar clinical features to MDD, such as social withdrawal and anhedonia (*9*); and by human studies demonstrating the depressogenic effects of pro-inflammatory therapies such as interferon (*10*).

Immune biomarker research in psychiatry has largely focused on the analysis of proteins in peripheral blood, which may reflect the activation state of multiple cell types both within and outside the circulation. Investigation of adaptive immunity and detailed immune cell phenotyping in depression have been less extensive (*11–17*). These parameters are important given recent genome-wide association studies in MDD implicating variants in major histocompatibility locus (MHC) genes that encode human leucocyte antigens (HLA), a group of glycoproteins that play a critical role in activating or suppressing adaptive immune cells (*18*). To date, most studies examining circulating immune cell subsets have been small, used limited phenotyping panels, and have generated somewhat conflicting findings.

Depression has been reproducibly associated with a leucocytosis, a raised neutrophil to lymphocyte ratio, and an increased CD4/CD8 T lymphocyte ratio (*19–21*), but studies have generated opposing observations on many other immune cell subsets including regulatory T cells, Th17 cells, NK cells and B cells (*11–15, 17*). These disparate findings may reflect differences in demographic variables – for example, sex, body mass index (BMI), tobacco use, and recent infection – that are associated with MDD and are known to impact immune cell numbers. In addition, studies have typically assessed the frequency of each immune cell subset relative to the total peripheral blood mononuclear cell (PBMC) pool, introducing a further challenge, as a relative decrease in the percentage of any given subset may reflect either a true decrease in cell number or a relative increase in other cell subsets.

Finally, it has become increasingly obvious that the syndromal diagnosis of MDD subsumes a heterogeneous patient population. Epidemiological data suggest at least two major subtypes of MDD; patients with ‘typical’ clinical features, such as reduced appetite and insomnia, who have normal circulating CRP and IL-6, and those with ‘atypical’ features, such as increased appetite, increased BMI and hypersomnia, who have evidence of immune activation (*22*). This highlights the need to accurately characterise and understand immune dysfunction in patients with MDD in order to identify patients who may benefit from targeted immunomodulation.

Here, we performed comprehensive immunophenotyping and deep clinical and demographic phenotyping in n=206 patients with MDD and n=77 controls, including circulating inflammatory proteins (CRP and IL-6) and peripheral blood cell counts. The MDD group was intended to be inclusive of a wide range of current symptom severity and anti-depressant drug treatment exposure and response, in order to support analysis of associations between immunological variables and multiple dimensions of clinical heterogeneity.

The prior hypothesis from the literature is that there is an ‘inflamed’ subgroup of MDD patients, who have evidence of immune activation and might be expected to respond better to immunomodulatory treatments (*22, 23*). Therefore, we sought to divide MDD participants into two groups – an inflamed and an uninflamed group – based on their immunophenotypic profile. We further hypothesized that patients with inflamed depression might not be a homogeneous group, and that inflamed depression may include patients with variable and differing underlying immune abnormalities. We thus used data-driven clustering to test whether subgroups of MDD patients demonstrated distinct immune phenotypes.

## Results

### Sample characteristics

We measured peripheral blood cell counts in participants with depressive disorder and non-depressed, age- and sex-matched healthy volunteers, recruited from five UK centers to the Biomarkers in Depression (BioDep) study (Chamberlain et al., 2018). Observer-rated and self-report questionnaires were used to determine clinical symptoms of depression and anxiety, as well as childhood and recent stress exposure: the Hamilton Depression Rating scale (HAM-D), the Beck Depression Inventory (BDI), the Chalder Fatigue Scale (CFQ11), the Snaith-Hamilton Pleasure Scale (SHAPS), the State-Trait Anxiety Inventory (STAI), the Childhood Trauma Questionnaire (CTQ) and the Life Events Questionnaire (LEQ), see **Supplementary Methods**. Absolute counts of 14 cell types were measured in all participants. Data passing quality control criteria were available on a sample comprising 206 MDD patients (143 female, 66 male) and 77 healthy volunteers (52 female, 25 male).

The MDD cases were clinically heterogeneous, comprising patients currently in remission as well as those who were severely depressed. Patients also varied in terms of current and historical antidepressant medication use (**Figure S1**). Case and control groups did not differ in terms of age, sex, or current use of tobacco or cannabis. As expected, the MDD group was significantly more depressed, anxious and fatigued than the non-depressed group. MDD patients also reported significantly more current stress, childhood trauma, alcohol use and unemployment than the healthy volunteers (**Table 1**).

**Table 1:** Demographic and clinical characteristics of the study population P-values for comparison of control vs. MDD by Mann-Whitney (MWU, continuous variables) or testing (categorical variables): *p<0.05, **p<0.01, and ***p<0.001. MDD, major depressive disorder; IQR, interquartile range; BMI, body mass index; STAI, Stait-Trait Anxiety Inventory. Number of missing data values for each variable are also shown (total n=283 participants).

| Participant characteristics | Control<br>(median,<br>IQR) | MDD<br>(median,<br>IQR) | P-value<br>(MWU) | Effect<br>size<br>$Z/\sqrt{n}$ | n (missing<br>data) |
| --- | --- | --- | --- | --- | --- |
| Number of participants | N = 77 | N = 206 | - |  |  |
| Age | 32.5<br>(28.3, 39.1) | 35.3<br>(28.7, 42.9) | 0.09 | 0.10 | 0 |
| BMI | 23.5<br>(21.3, 27.6) | 26.6<br>(23.0, 31.2) | ***0.0008 | 0.20 | 8 |
| Hamilton depression rating | 0<br>(0.0, 1.0) | 17<br>(14.0, 20.0) | ***2E-53 | 0.74 | 0 |
| Beck depression inventory | 1<br>(0.0, 3.0) | 24<br>(15.0, 31.2) | ***8E-43 | 0.70 | 6 |
| Chalder fatigue score | 11<br>(8.0, 11.0) | 19<br>(14.0, 23.5) | ***4E-33 | 0.64 | 3 |
| Snaith-Hamilton Pleasure<br>Scale | 0<br>(0, 0) | 4<br>(1, 7) | ***2E-24 | 0.58 | 4 |
| STAI (state subscale) | 25<br>(22.0, 29.0) | 50<br>(38.5, 57.5) | ***2E-39 | 0.68 | 3 |
| STAI (trait subscale) | 27<br>(24.0, 32.0) | 60<br>(52.0, 68.0) | ***1E-48 | 0.73 | 3 |
| Childhood trauma score | 35.0<br>(33.0, 38.5) | 49.5<br>(40.0, 62.0) | ***3E-19 | 0.51 | 6 |
| Recent stressors (z-score) | -0.9<br>(-0.9, -0.3) | -0.2<br>(-0.4, 0.5) | ***1E-10 | 0.37 | 4 |
| Number of previous<br>ineffective antidepressant<br>treatments (<75% response) | - | 1.0<br>(1.0, 3.0) | - | - | 7 |
| | Control<br>(percent) | MDD<br>(percent) | P-value | $\chi^2$ | n (missing<br>data) |
| Female sex | 68% | 69% | 0.8 | 0.09 | 0 |
| Unemployed (including for<br>medical reasons) | 0% | 23% | ***0.0005 | 21.4 | 3 |
| Current tobacco use | 11% | 12% | 0.8 | 0.09 | 4 |
| Current alcohol use | 33% | 48% | *0.03 | 4.81 | 5 |
| Current cannabis use | 3% | 6% | 0.3 | 1.57 | 5 |
| Current antidepressant use | - | 75% | - |  | 5 |

### Case-control univariate analysis: peripheral blood cell counts and inflammatory proteins

We hypothesized that MDD patients would show increased peripheral cell counts and inflammatory proteins compared to healthy controls. Cell counts were estimated by combining the percentage count data from flow cytometry with absolute counts from a standard hematology panel; see **Methods** and **Table S2**. Using multiple univariate case-control comparisons, there was some evidence for abnormality of three immune cell counts and two inflammatory proteins in MDD cases compared to controls (**Figure 1A, Table S3**). Serum CRP and plasma IL-6 concentrations, as well as absolute counts of neutrophils, intermediate monocytes and CD4^+^ (helper) T cells, were all increased on average in the MDD group (*P* < 0.05). When the comparison between two groups was corrected for the multiple biomarkers tested (16; FDR *P* < 0.05), only CRP and CD4^+^ T cells remained significantly different between the groups.

**Figure 1:**
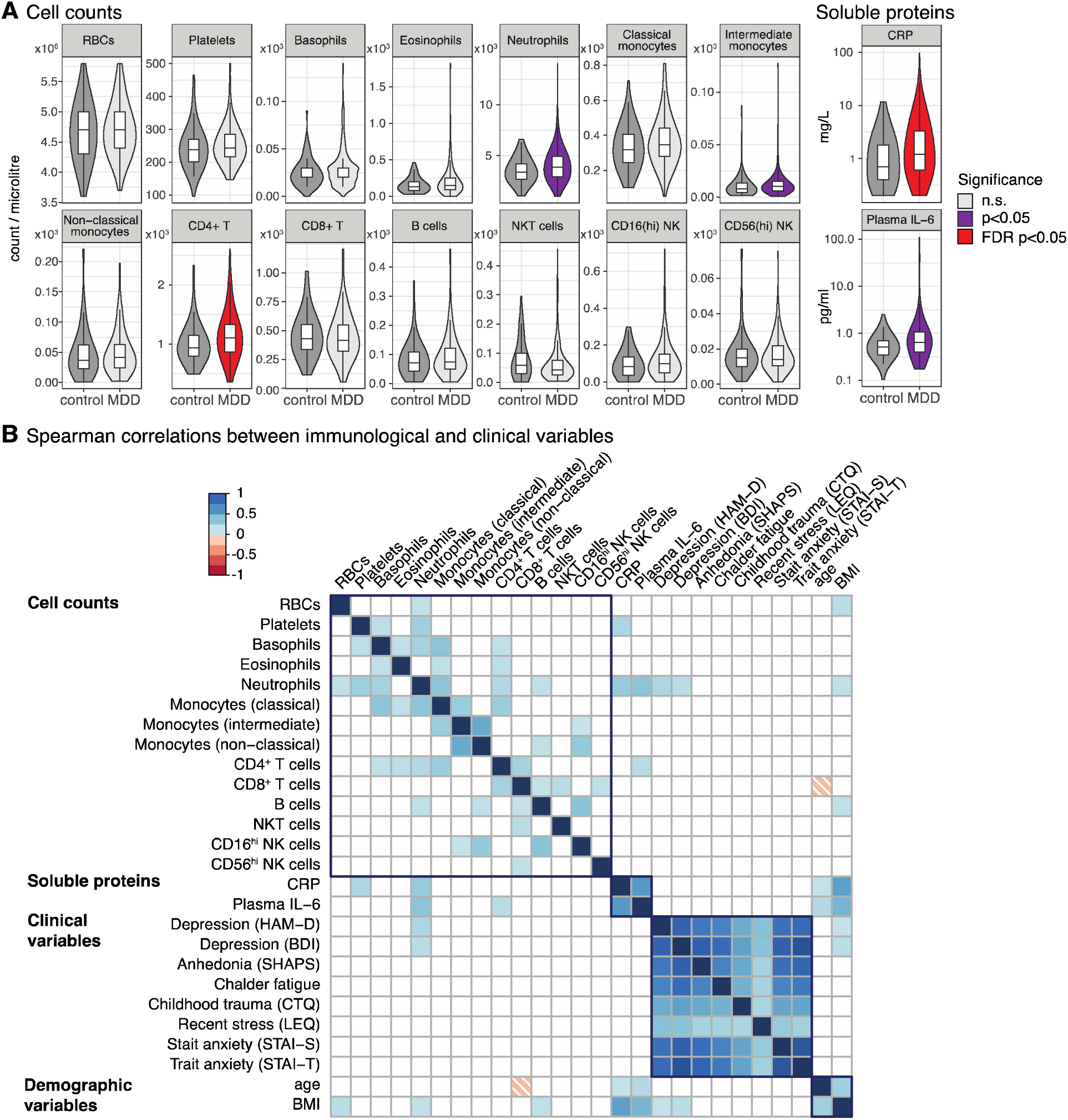
Peripheral immunophenotypes in MDD and control participants. **(A)** Comparison of 14 absolute cell counts, high sensitivity CRP and plasma IL-6 in major depressive disorder (MDD, n=206) and matched controls (n=77). Boxplots show median and interquartile range, with the outer violin shape showing the full distribution of data. Color indicates statistical significance by Mann Whitney U test (purple *P* < 0.05; red FDR *P* < 0.05). FDR p-values are corrected for 16 multiple comparisons. See **Table S3** for underlying data and effects sizes. **(B)** Spearman correlations between immunological, clinical and demographic variables. Only those correlations significant at *P* < 0.001 are shown. Color indicates the correlation coefficient (Spearman’s ρ); dark blue outlines group together similar variables.

### Case-control multivariate analysis: principal components of correlated cellular, protein and clinical variables

We estimated correlations between all immunological, clinical and demographic variables in the entire sample set (N=283) (**Figure 1B**). Immune cell counts and inflammatory protein concentrations were generally correlated with each other, as were questionnaire measures of symptom severity and stress. The only immunological variable significantly correlated with clinical symptom scores was the neutrophil count, which correlated positively with HAM-D score (Spearman’s ρ = 0.27, *P* < 0.00001) and BDI score (ρ = 0.25, *P* = 0.00003). Correlations between these variables within the MDD group only were similar to those in the dataset as a whole (**Figure S3**).

Given the highly correlated cell counts, we used principal components analysis (PCA) to define two orthogonal, linear combinations of the 14 cell subset counts which accounted for the greatest proportion of the variance-covariance (**Figure 2A**). Each component can be regarded as the expression of a system of developmentally or functionally related cell subsets. The first principal component (PC1), which accounted for 19% of the total variance-covariance, was a weighted average of all cell counts, most strongly weighted on myeloid cells (especially neutrophils, basophils and classical monocytes) and CD4^+^ T cells (**Figure 2A**). The second principal component (PC2) accounted for 10% of the total variance-covariance and represented a combination of increased non-classical monocytes and CD16^hi^ NK cells and decreased classical monocytes (**Figure S4A**). The structure of PC1 was similar when PCA was performed separately for each sex (**Figure S4**).

**Figure 2:**
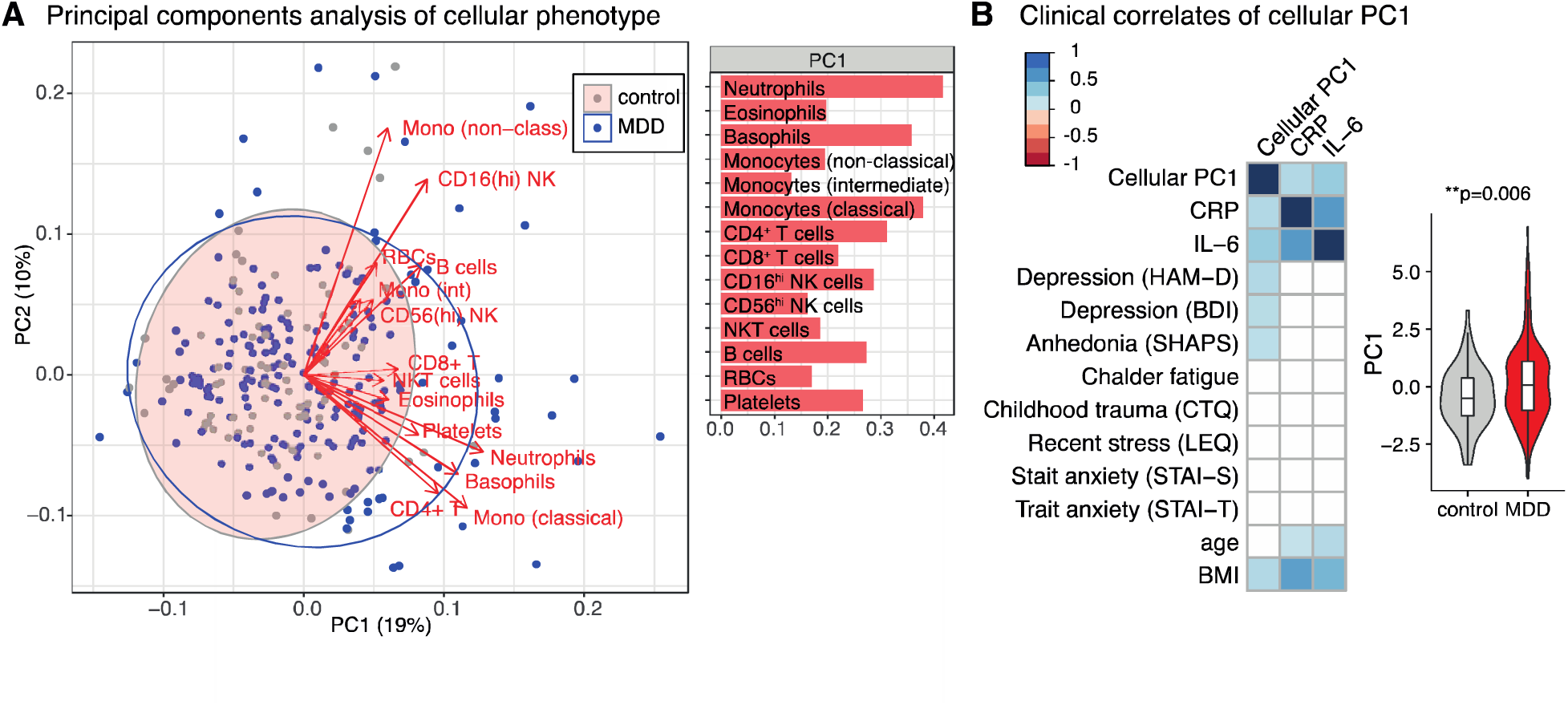
Principal components analysis of cellular immunophenotypes. **(A)** Principal components analysis (PCA) for the 14 absolute cell counts across all participants (n=283). On the left panel, each point (MDD in blue; controls in grey) represents one participant’s scores on the first two principal components (PC1 and PC2). Red arrows show the loadings of each cell count on the first two principal components. Ellipses show the 95% confidence ellipse for each group. Right hand panel shows the PCA eigenvector for PC1. See **Figure S4** for PC2 and PCA excluding patients with minor inflammatory conditions, and for each sex separately. **(B)** Spearman correlations between the first principal component of the cellular immunophenotype (PC1), clinical features, demographic features, and peripheral proteins (n=283 participants). Only correlations significant at *P* < 0.001 are shown. Color indicates the correlation coefficient (Spearman’s ρ). The right-hand side boxplot shows the PC1 scores for MDD cases and controls (Mann-Whitney test, estimate=0.54, effect size=0.16, *P* = 0.006). Boxplots show median and interquartile range, outer violin shape shows the full distribution of data.

The extent to which each participant’s immune cell profile corresponded to the first PC was summarized by their PC1 score. Participants with high positive PC1 scores had generally increased counts, especially in myeloid/CD4^+^ subsets; whereas low negative PC1 scores indicated decreased counts across all cell subsets. PC1 scores were positively correlated with serum CRP (ρ = 0.26, *P* < 0.00001), and IL-6 (ρ = 0.34, *P* < 0.000001) concentrations, indicating that patients with elevated peripheral inflammatory markers have increased counts of both innate (neutrophil) and adaptive (CD4+) immune cell subsets. The MDD cases on average had higher scores than non-depressed controls on expression of this inflammatory cell system (Mann Whitney U: *P* = 0.006, standard effect size, *r* = 0.16) (**Figure 2B**). Observer-rated depressive symptoms (HAM-D, ρ = 0.26, *P* < 0.00001), self-reported depressive symptoms (BDI, ρ = 0.24, *P* = 0.00005), anhedonia (SHAPS, ρ = 0.23, *P* = 0.0001), and BMI (ρ = 0.24, *P* < 0.001) were all positively correlated with PC1 scores (**Figure 2B**).

A scatterplot of participants in the space of the first two PCs (**Figure 2A**) shows that most depressed cases have PC scores overlapping with the distribution of PC scores in the healthy controls, indicating that the majority of depressed patients have a healthy cellular phenotype. However, there is a subgroup of MDD patients with more extreme scores, indicating increased cell numbers, mainly in the myeloid compartment.

### Case-control multivariate analysis: PLS discriminant analysis of immune cell phenotypes

To test the multivariate hypothesis that a combination of correlated immune cell counts was predictive of the case-control difference between groups, we used partial least squares (PLS) discriminant analysis (**Figure 3A and S4E**). This demonstrated that the two groups could be significantly discriminated from each other by a discriminant function of the 14 cell counts (*P* = 0.002, permutation test) although this only accounted for 6.3% of the variability in diagnostic status. Absolute cell counts for CD4^+^ T cells, neutrophils and eosinophils were significantly weighted on the discriminant function, indicating that a combination of these cell counts (not simply each cell count individually) was most predictive of classification into MDD versus control groups.

**Figure 3:**
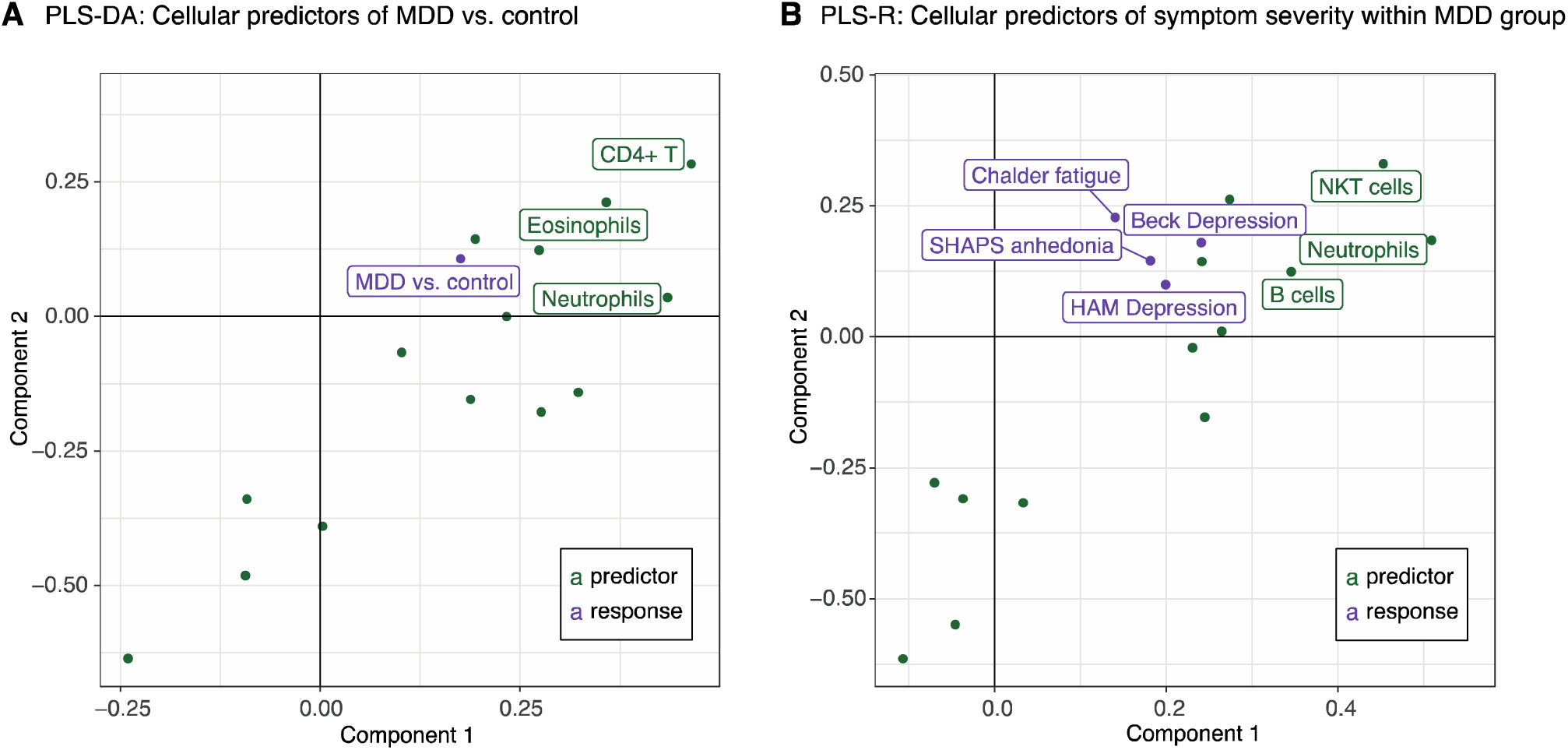
Cellular predictors of MDD status and symptoms severity. **(A)** Partial least squares discriminant analysis (PLS-DA) for the predictors of case-control status. Response variable is major depressive disorder (MDD)/control status (purple point), predictor variables are the 14 absolute cell counts from **Figure 1A** (green points). Analysis includes all participants (n=283). A single component PLS model (Component 1) is significantly predictive of MDD status by permutation testing (*P* = 0.002). Of the 14 cell types, only those with significant weights in the model are labelled: neutrophils, eosinophils and CD4^+^ T cells (bootstrapped Z-score >3, see **Methods**). See also **Figure S4E**. **(B)** Partial least squares regression (PLS-R) for the predictors of depressive symptom severity within the MDD group. Response variable is the matrix of symptoms scores (shown in purple), predictor variables are the 14 absolute cell counts (green points). A single component PLS model (Component 1) is significantly predictive of MDD severity by permutation testing (*P* = 0.001). Of the 14 predictor cell types, only those with significant weights in the model are labelled: neutrophils, NKT cells and B cells (bootstrapped Z-score >3, see **Methods**). Analysis includes MDD cases only (n=199 with full clinical scores available). SHAPS = Snaith-Hamilton Pleasure Scale. HAM = Hamilton Depression Rating Scale.

### Within-group analysis: PLS regression of immune cell counts associated with severity of depression

We used PLS regression to test the hypothesis that a linear combination of immune cell counts predicted MDD symptom severity within the MDD group. Using a matrix of clinical symptom scales as the multivariate response variable (HAM-D, BDI, Chalder Fatigue and SHAPS), we found that a single PLS component was significantly associated with the severity of depressive symptoms (*P* = 0.001, permutation test) and explained 7.3% of the variance in symptom severity scores. The cell counts significantly weighted on the PLS component were neutrophils, NKT cells and B cells, indicating that a combination of these three cell counts was most strongly related to symptom severity, especially as self-reported by the BDI (**Figure 3B**).

### Within-group analysis: forced two-way clustering of MDD cellular immunophenotypes

These results indicate significant immunological differences between MDD patients and controls, but also considerable heterogeneity of immune cell phenotypes within the MDD group. We thus investigated the hypothesis that there were subgroups or clusters within the MDD group (N=206), characterized by distinct immune profiles.

To identify a stable two-way partition of MDD cases, we used Gaussian finite mixture modeling of 90% subsamples of the cellular dataset followed by consensus clustering across the partitions generated (see **Methods**). This analysis, based only on cell count data, identified a cluster comprising 39% of MDD cases (‘inflamed depression’, n=81) who had broadly elevated immune cell counts and inflammatory proteins compared to the second cluster of patients (‘uninflamed depression’, n=125) (**Figure 4A**). Compared to uninflamed cases, inflamed MDD cases had higher numbers of all monocyte subsets, all granulocyte subsets, CD16^hi^ NK cells, NKT cells, B cells, T cells and platelets (**Figure 4B**). In terms of their clinical phenotype (**Figure 4D**, **Table S4**), inflamed cases had a significantly higher severity of observer-rated depressive symptoms (HAM-D, *P* = 0.0002) and self-reported depressive symptoms (BDI, *P* = 0.01). Cases in the inflamed group had a median HAM-D score of 19, corresponding to moderate severity depression, while uninflamed cases had a median HAM-D score of 16, corresponding to mild severity depression (*24*). Inflamed vs. uninflamed cases also had double the rate of unemployment (33% vs.17%, *P* = 0.008) and higher blood concentrations of CRP, plasma IL-6 and triglycerides (**Figure 4C**); but inflamed cases did not have higher BMI or higher symptom scores on clinical measures of ‘typical’ depression (**Figure S5**).

**Figure 4:**
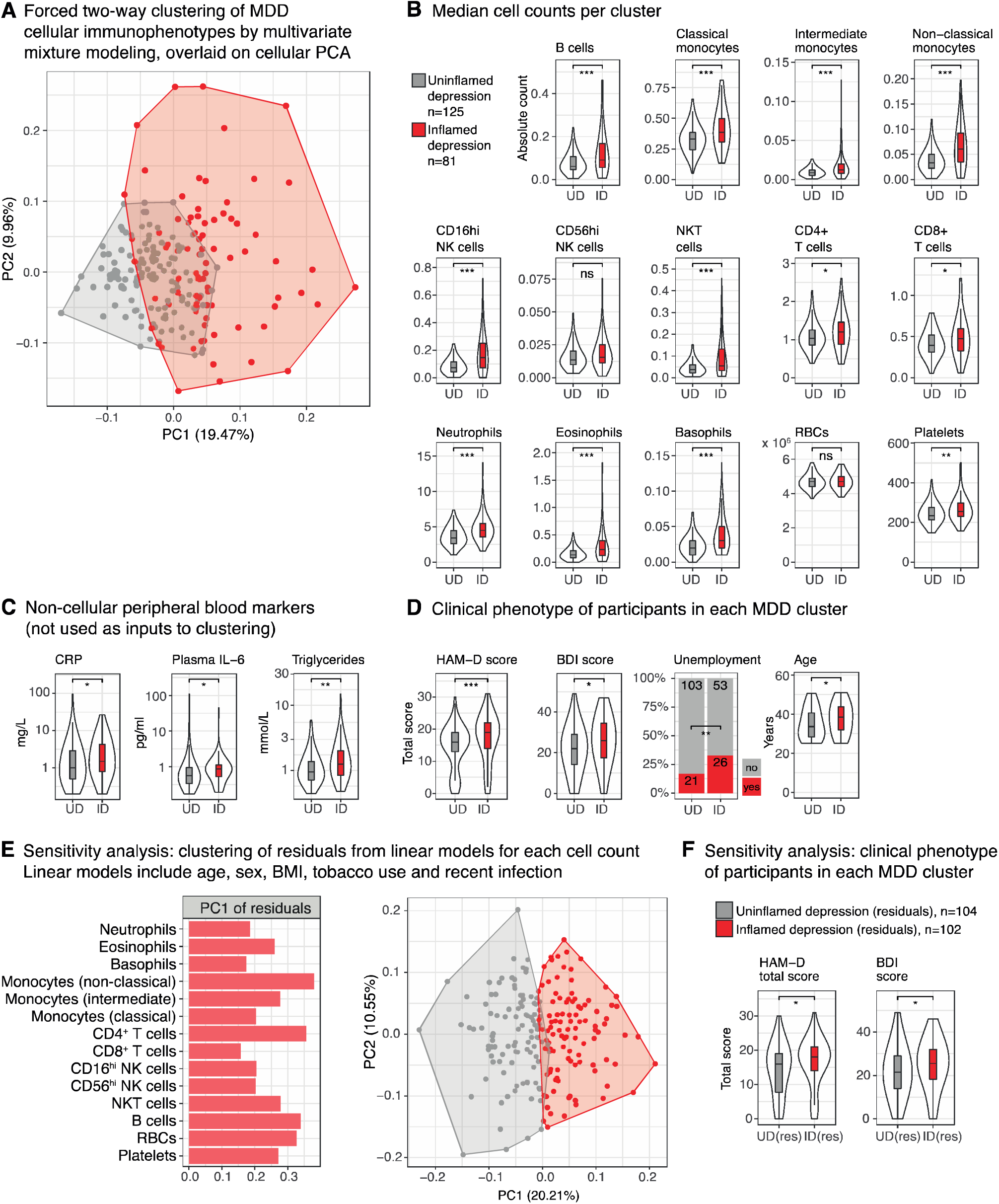
Forced two-way clustering of MDD cellular immunophenotypes. **(A)** Gaussian finite mixture modelling of the cellular phenotypes for MDD cases (n=206). Forced two-way mixture modelling identified two clustered immunophenotypes, uninflamed depression (UD, n=125 cases, grey) and inflamed depression (ID, n=81, red). Plot shows the PCA scores for each case on cellular PC1 and PC2 (**Figure S5A**), with cluster membership indicated by color. **(B, C, D)** Comparisons between the two clusters. Boxplots show median and interquartile range for each cluster, with the outer violin shape showing the full distribution of data. Effects of cluster were tested by Mann-Whitney U or (for unemployment), \**P*<0.05, \*\**P*<0.01, \*\*\**P*<0.01. **(B)** Absolute cell counts (inputs to clustering). **(C)** Peripheral blood markers: C-reactive protein (CRP), interleukin-6 (IL-6) and triglycerides (not used as inputs to clustering). **(D)** Clinical phenotype of participants in each cluster. Bar annotations indicate participant numbers. See **Table S4** and **Figure S5** for statistics, further clinical and demographic measures and item-level scores corresponding to MDD ‘typicality’. **(D)** Sensitivity analysis: principal components analysis (PCA) of the 14 residual cell counts following linear modelling to regress out the effects of body mass index (BMI), age, sex, current tobacco use and recent infection on each cell count (MDD cases only, n=206). Left-hand panel shows the eigenvector for the first principal component of the residual cell counts (PC1). Right hand panel shows the results of forced two-way clustering of the residual cell counts, which identified two immunophenotypes, uninflamed depression (n=104, grey) and inflamed depression (n=102, red), overlaid on a scatterplot of PCA scores. Inflamed depression is characterized by higher residual cell counts across all cell types (**Figure S6A**). **(E)** Depression rating scores for cases in each cluster in the sensitivity analysis. Effects of cluster were tested by Mann-Whitney U, \**P*<0.05, \*\**P*<0.01. Other clinical features and demographic variables did not differ significantly between the clusters. HAM-D = Hamilton Depression Rating Scale (practitioner-administered), BDI = Beck Depression Inventory (self-report).

Inflamed cases were slightly older than uninflamed cases (median age 38 years vs. 34 years) and more likely to be smokers (19% vs. 7%). Otherwise, there were no significant differences in terms of potential confounding factors including sex, study center, current antidepressant use, alcohol use, cannabis use, reported recent infection, or presence of minor inflammatory disease (**Figure S5B**, **Table S4**).

### Within-group analysis: Data-driven clustering of MDD cellular immunophenotypes

Given the heterogeneity in immunophenotypes among the depressed cases, we hypothesized that there may be more subtle, stratified immune cell phenotypes beyond a simple inflamed vs. uninflamed binarization. We therefore took an unbiased, data-driven approach to identifying subgroups of cases associated with a distinctive peripheral blood cell profile, without specifying the expected number of subgroups, in contrast to the prior analysis where we pre-specified two possible subgroups of cases: inflamed or uninflamed depression. To identify the stable and distinct immunophenotypes present in MDD cases, we used Gaussian finite multivariate mixture modeling of 90% subsamples of the data to generate clusters, selecting the optimal cluster number for each partition by BIC. We then performed consensus clustering of the partitions generated, which identified 4 subgroups or clusters of cases (**Figure 5, Figure S7, Table S5**).

**Figure 5:**
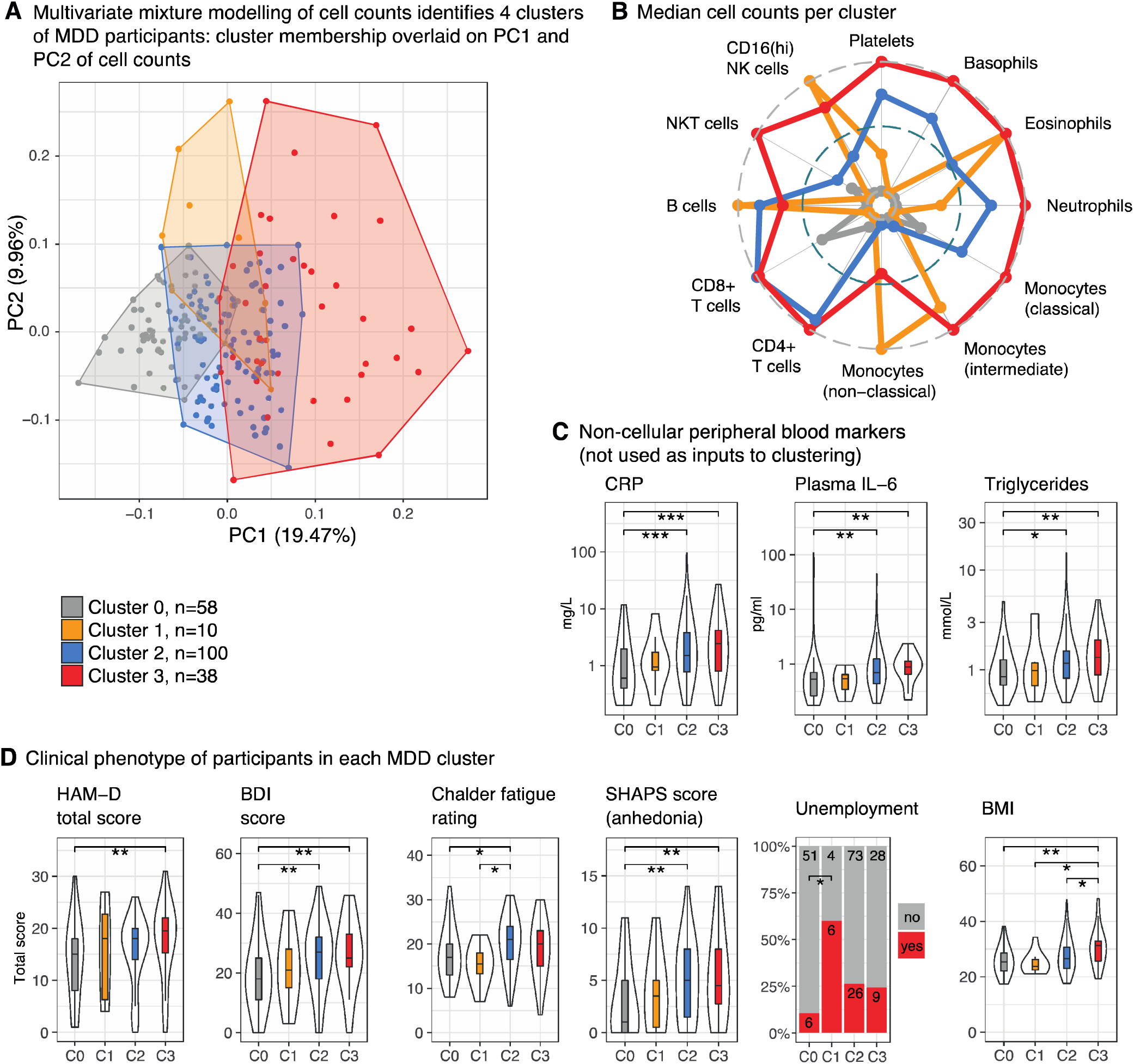
Stratified MDD cellular immunophenotypes. **(A)** Data-driven Gaussian finite mixture modelling of the cellular phenotypes for MDD cases (n=206) identified four discrete clusters (immunophenotypes). Plot shows the PCA scores for each participant on cellular PC1 and PC2, with cluster indicated by color. The arbitrary cluster numbers and colors are used consistently throughout this figure to designate each cluster (cluster 0, grey, n=58 cases; cluster 1, orange, n=10; cluster 2, blue, n=100; cluster 3, red, n=38). **(B)** A radar plot shows the characteristic cellular immunophenotype of each cluster. Points represents the median value of the 14 absolute cell counts for each of the four clusters, rescaled onto a 0 to 1 range (with higher values on the outside of the plot) to highlight relative differences between clusters. Cluster differences are significant for all counts shown (Kruskal-Wallis *P*<0.05); red blood cell and CD56^hi^ NK cell counts did not differ between the clusters and are not shown. See **Figure S7A** for underlying data and statistics. **(C)** Peripheral blood markers and **(D)** clinical and demographic features for each cluster. Boxplots show the median and inter-quartile range of the relevant variable for each cluster, violin outline shows the full distribution of data. The effect of cluster on each continuous feature is tested by Kruskal-Wallis testing. Where *P* < 0.05 for the overall Kruskal-Wallis test, we performed post-hoc Conover tests to identify which pairs of clusters differ for that feature – for these variables, each cluster was compared to every other cluster. For unemployment, the bar chart indicates the percentage of participants in each cluster and bar annotations indicate participant numbers. Clusters were compared by testing, with post-hoc tests to compare pairs of clusters. All p-values shown are corrected for the 6 pairwise cluster-cluster comparisons performed: FDR *P* *<0.05, **<0.01, and ***<0.001. Pairwise comparisons which were non-significant following FDR correction are not shown. For further statistics, demographics and item-level scores corresponding to MDD ‘typicality’ see **Figures S7, Table S5**. HAM-D = Hamilton Depression Rating Scale (practitioner-administered), BDI = Beck Depression Inventory (self-report), SHAPS = Snaith-Hamilton Pleasure Scale.

Each of the 4 subgroups or clusters of cases was characterized by a distinct immune cell profile (**Figure 5B, Figure S7A**). One cluster comprised 58 cases (28%) with low counts for all cell subsets and low levels of CRP and plasma IL-6 – subsequently referred to as the uninflamed subgroup (C0). The other three clusters (C1-3) showed evidence of immune activation as follows. Cluster 1 (C1) comprised only 10 cases (5%) and was characterized by selective increases in B cells, non-classical monocytes, intermediate monocytes and CD16^hi^ NK cells, with no accompanying increase in classical monocytes or neutrophils relative to uninflamed patients. Cluster 2 (C2, 100 patients, 49%) and Cluster 3 (C3, 38 patients, 18%) shared a common increase in classical monocytes, neutrophils, eosinophils, basophils, B cells, CD4^+^ T cells and platelets compared to uninflamed patients. Both of these clusters were associated with increased CRP, IL-6 and triglycerides (**Figure 5C**) and we refer to these as the inflamed depression subgroups. Comparing these two inflamed subgroups, C3 had a stronger myeloid bias compared to C2, with significantly higher numbers of classical monocytes, intermediate monocytes, non-classical monocytes and neutrophils. Overall, adaptive immune cells were relatively more prominent than myeloid cells in the immunophenotype of C2 compared with that of C3 (**Figure 5B**). This four-way data-driven stratification of cases was not simply nested within the forced two-way partition into inflamed and uninflamed depression. All uninflamed cases in the data-driven four-subgroup classification were also classified as uninflamed in the two-group classification; and all cases the myeloid-biased inflamed subgroup in the four-group analysis (C3) were binarized as inflamed in the two-group analysis. However, cases in one of the other inflamed sub-groups in the four-group analysis (C2, adaptive-biased) were assigned to both uninflamed and inflamed subgroups by the binary analysis (**Table S6**).

We next tested whether the four immune cell count-defined subgroups of cases had different clinical phenotypes (**Figures 5D, Table S5**). Subgroup classification had a significant effect on observer-rated depressive symptoms (HAM-D, *P* = 0.009), self-reported depressive symptoms (BDI, *P* = 0.002), anhedonia (SHAPS, *P* = 0.003), fatigue (CFQ11, *P* = 0.005) and unemployment (*P* = 0.004). On pair-wise post-hoc testing, both inflamed subgroups (C2 and C3) were associated with increased depression severity compared to uninflamed depression. C3 (inflamed, myeloid-biased) cases vs. C0 (uninflamed) cases had significantly higher severity of observer-rated depressive symptoms (HAM-D, FDR *P* = 0.004), self-reported depressive symptoms (BDI, FDR *P* = 0.006) and anhedonia (SHAPS, FDR *P* = 0.006). C2 (inflamed, adaptive-14 biased) cases vs C0 cases likewise had a significantly higher severity of self-reported depressive symptoms (BDI; FDR *P* = 0.003), anhedonia (SHAPS; FDR *P* = 0.004) and fatigue ratings (CFS; FDR *P* = 0.02). Median HAM-D scores in the inflamed subgroups C3 and C2 were 19.5 and 18 respectively (i.e. moderate depression) compared to 15 for uninflamed C0 cases.

Immune cell count-derived subgroups of cases were not significantly different from each other in terms of anxiety, anti-depressant treatment-resistance, typical or atypical depression symptom scores, recent stress, or childhood trauma (**Figure S7**). C3 inflamed cases had higher median BMI than uninflamed cases (FDR *P* = 0.001), but C2 inflamed cases did not. C1 contained few cases (n=10), but nonetheless was significantly associated with unemployment (60% vs. 11% for C0 cases, FDR *P* = 0.006).

### Sensitivity analyses demonstrate inflammatory cellular phenotypes in depression independent of BMI, age, sex, recent infection and tobacco use

For the 4-subgroup classification of cases based on immune cell counts, we investigated the potentially confounding effects of other variables. There was no association between classification and sex, recent infection, presence of minor inflammatory disease, current tobacco, alcohol or cannabis use, or current antidepressant treatment (**Figure S7B**, **Table S5**). The effect of age was overall significant (*P* = 0.02) but none of the post-hoc pair-wise comparisons were significant. There was an association between study center and subgroup (*P* = 0.04), driven by the smallest subgroup (C1) only comprising cases from two of the five centers (**Figure S7B**).

In order to further explore the effect of demographic variables on MDD immunophenotypes, we used linear regression to “factor out” the effects of age, sex, BMI, recent infection history and tobacco use before analysis of residualised counts of the 14 immune cell subsets. We found that the first principal component of the residual cell counts was essentially the same as that for the absolute counts, with PC1 again representing a weighted average of increases in all 14 cell counts (**Figure 4E**), but with neutrophils and basophil counts now less prominent and non-classical monocytes more prominent contributors to PC1, compared to the unadjusted analysis (**Figure S5A**). Recapitulating the results for the absolute cell counts, PLS-R identified a single component model as predictive of MDD severity (permutation test, *P* = 0.01), with neutrophil and NKT residual cell counts significantly weighted on this component. We next performed a forced two-way clustering analysis of MDD cases as above, but using residual cell counts rather than absolute counts as inputs to the clustering (**Figure 4E**). This analysis again identified an inflamed subgroup of patients with higher residual counts across all 14 cell types compared to uninflamed patients (**Figure S6A**). Cases in the inflamed subgroup had increased HAM-D and BDI scores (**Figure 4F**) compared with cases in the uninflamed subgroup, as well as increased triglycerides (**Figure S6B**). There were no significant differences between the subgroups in CRP, IL-6 or other demographic or clinical variables. This analysis demonstrates clear associations between depressive symptoms and stratified MDD immunophenotypes independent of age, sex, BMI, infection and tobacco use.

## Discussion

### Immune cell counts as markers of inflamed depression

Previous studies have suggested that some patients with MDD have elevated pro-inflammatory cytokines and CRP in peripheral blood and abnormalities of circulating immune cells (*3, 6, 7, 19*). Here, we confirm that, on average, patients with MDD have increased myeloid cells, lymphoid cells and inflammatory proteins compared to healthy controls (**Figure 1A**). We further demonstrate that these correlated immunological changes are associated with symptom severity (**Figure 2, 3B**). However, not all MDD patients had evidence of inflammation, consistent with previous reports (*22, 23*). We find that MDD patients can be divided into inflamed and uninflamed groups based on clustering of their cellular immune profiles, with patients in the inflamed subgroup showing greater depression severity (**Figure 4**). Notably, our data-driven approach revealed that inflamed depression is not a monolithic phenotype, but rather, comprised distinct immunophenotypic subgroups, that differed in terms of relative increases in innate or adaptive immune cells (**Figure 5**). One inflamed subgroup showed a myeloid-dominant leukocytosis and the other a relative lymphoid bias. Notably, these two inflamed immunophenotypes could not be distinguished from each other by CRP or IL-6 levels, or by symptom scores. These data implicate both innate and adaptive immunity in the pathogenesis of inflamed depression, but their relative contribution may differ in different patients. By increasing the granularity of immunophenotyping, we have identified previously unappreciated heterogeneity within the inflamed depression group, potentially reflecting MDD subgroups with distinct pathogenesis.

One obvious question raised by our analysis is the relationship between the different inflamed subgroups identified. We do not know whether these immunophenotypes are stable over time in individual patients, reflecting distinct underlying mechanisms, or whether patients might move between one subgroup and another, perhaps as disease progresses. It may be, for example, that the subgroups represent a continuum over time, where patients start with a phenotype dominated by innate immune activation, and subsequently develop more pronounced activation of the adaptive immune system. Future longitudinal studies will begin to address these questions.

Most work in human depression and animal models of mood disorder has focused on the innate immune system. MDD-associated changes in peripheral proteins such as IL-6 are typically interpreted as representing overactivity of myeloid cells, but there is now increasing recognition of the potential importance of adaptive immunity to mood disorder (*25*). Indeed, adaptive cells, particularly B cells, can act as a major source of IL-6 in the context of immunological challenge (*26*). A role for adaptive immunity is supported by the association of MDD with variants in the MHC region (*18*) which encodes genes for human leucocyte antigens (HLA), glycoproteins critical for adaptive immunity. HLA antigens expressed by antigen presenting cells, including B cells, are required for CD4^+^ T cell activation. Our data show evidence of variable adaptive immune activation in both inflamed MDD subgroups (**Figure 5**).

### Neutrophil abnormalities in MDD

When the cell counts were considered individually, only neutrophils correlated with depressive symptom scores (**Figure 1B**). Neutrophil count predicted MDD vs. control status in a discriminant analysis (**Figure 3A**), predicted symptom severity within the MDD group (**Figure 3B**), and was significantly higher in the inflamed MDD subgroups with increased depressive symptoms (**Figures 4, 5**).

Clinical studies have previously described a univariate association between neutrophils and depression (*19–21, 27*). Neutrophilia is usually seen in the context of bacterial infection as part of ‘emergency granulopoiesis’, where endothelial G-CSF leads to increases in both the production and release of neutrophils (*28*). Elevated G-CSF has been observed in patients with MDD (*5*) and glucocorticoids increase both G-CSF and neutrophil numbers (*29*). The hypercortisolemia observed in some MDD patients (*30*) may thus underpin the relative neutrophilia observed in MDD patients in our study.

How might the changes in neutrophils and CD4^+^ T cells be related? There is an increasing appreciation that neutrophils may play a role in adaptive immunity, including the demonstration of antigen presentation to CD4^+^ T lymphocytes via major histocompatibility complex II (MHCII) in vitro (*31, 32*). CD4^+^ T cells may in turn impact neutrophils. This has been best described in the context of tissue inflammation mediated by Th17 cells, where IL-17 secretion promotes epithelial cell secretion of neutrophil recruiting chemokines and survival factors (*33*).

### Potential sources of cellular immune abnormalities in MDD

Stress and infection are risk factors for subsequent depression (*34, 35*). Both can also cause hematopoietic stem cell proliferation and consequent increases in immune cell numbers in the peripheral blood (*36, 37*), supporting a model where inflammatory insults and psycho-social stresses cause release of innate immune cells into the circulation from reserves in the bone marrow and spleen (*38*). However, in our data, the inflamed subgroups of depressed cases were not significantly associated with reported recent stress or infection, nor were they associated with antidepressant treatment responsiveness. The factors which account for the observed immunophenotypes are thus unclear, but both genetic and environmental factors could play a role. Future studies incorporating detailed clinical phenotypes and comprehensive immunophenotyping, married to an analysis of genetic variants, will help to resolve these questions.

### Immune cell homeostasis in MDD

While we have a good understanding of many of the stimuli which increase circulating immune cell numbers, our understanding of how a normal range is restored or maintained is more limited (*39*). Whatever the cause, persistently raised peripheral blood cell counts in depression must reflect either an increased release from primary lymphoid organs, increased proliferation in secondary lymphoid organs or non-lymphoid tissues, increased immune cell lifespan, decreased extravasation of cells into tissues, or a combination of these factors. The lifespan of immune cells in the blood ranges from <1 day for neutrophils; 1-7 days for monocyte subsets; and weeks to years for B and T cells (*40*). The altered immune profiles we observe may be the result of an ongoing stimulus e.g. stimulation of pattern recognition receptors by damage- or pathogen-associated molecular patterns (DAMPs or PAMPs). Alternatively (or additionally), the immunophenotypes may reflect the memory of a now-terminated risk factor e.g. an infection or a stressful event. This could occur via epigenetic changes in the longer-lived bone marrow precursors (*41*), or remodeling of neural control of immune cell release from the bone marrow or spleen. Kinetic studies of immune cells in humans (*42*) may help identify how cellular kinetics are altered in MDD.

It remains unclear how these observed alterations in peripheral immune cells might affect mood. Candidate mechanisms suggested by animal models include a direct effect of cytokines infiltrating into the brain (*43*); afferent signalling of inflammation via cytokine receptors on vagal nerve afferent terminals; the induction of metabolic enzymes and transporters that decrease the synaptic availability of serotonin and increase the production of neurotoxic glutamatergic agonists (*44*); or trafficking of peripheral immune cells to the meningeal immune system where they can act locally to affect behavior (*45*). Neutrophils can traffic to the brain and neutrophil depletion has been shown to mitigate the effects of inflammation on behavior in animal models (*46*).

### The role of ‘confounds’ and co-morbidities in MDD immunophenotypes

BMI, age and sex are risk factors for depression and are also known to affect the peripheral immune system. However, there were no significant pairwise differences in age or sex between the four immune cell subset-defined subgroups, and only the myeloid-dominant inflamed subgroup (C3) was associated with increased BMI compared to the uninflamed cases (C0). Furthermore, sensitivity analyses using residual cell counts (where variability due to potentially confounding factors had been attenuated by prior regression) demonstrated that significantly greater depressive symptoms in the inflamed vs uninflamed subgroups was independent of subgroup differences in age, sex, BMI, recent infection and tobacco use. BMI-associated alterations in cell counts may be of relevance to understanding the cause of depressive symptoms: to discount them entirely as confounds seems over-simplistic at this stage.

There are no universally accepted subgroups of depression derived from clinical data, but a distinction akin to the contrast between ‘typical’ and ‘atypical’ has recently gained traction as biologically meaningful (*47*), with atypical (but not typical) depression associated with increased levels of CRP, IL-6 and triglycerides (*22*). In our data, patients in the inflamed subgroups showed these biochemical changes and these subgroups may overlap with (or be nested within) this symptom-derived group, but we did not have the necessary clinical information available to classify patients into an atypical depression group.

### Implications for whole-blood assays in MDD

Our demonstration of altered immune cell counts in depression has important implications for the interpretation of whole-blood assays. Peripheral blood cytokines and whole-blood gene expression measurements reflect the pooled effects of multiple cell types. Analysis of whole-blood data may thus appear to imply changes in genes and pathways which have not in fact been altered in any specific cell subset. Functional and transcriptional assays of sorted cell subsets, or analyses of whole blood gene expression controlling for the effects of differences in cell subset counts, will be crucial to furthering our understanding of transcriptional as well as cellular abnormalities in depression.

### Methodological limitations

Most MDD cases were recruited using a targeted strategy aiming to recruit participants equally to three groups: currently depressed and unmedicated; currently depressed and medicated with poor treatment response; and medicated with good treatment response. We did not include inpatients with MDD, who tend to have more severe symptoms. This will influence the generalizability of our results. Here, cell counts were derived by combining relative cytometry counts with absolute haematology counts: cytometry techniques using beads for absolute quantification may provide more accurate estimated counts.

### Implications for stratification

Our demonstration of distinct patterns of immune activation in MDD has implications for treatment strategies, indicating that stratification of therapies targeting different arms of the immune system may be required. It also informs the design of future clinical trials of immunomodulatory therapy in depression, highlighting the need for better immunophenotyping to allow treatment selection and to provide a deeper understanding of the cellular mechanisms of inflamed depression.

## Materials and Methods

### Study design

We measured peripheral blood cell counts in participants with current or past history of depression, and non-depressed, age- and sex-matched healthy volunteers, recruited from five UK centers to the Biomarkers in Depression (BioDep) study, a cross-sectional observational study (Chamberlain et al., 2018). Demographic and clinical characteristics of the sample are outlined in **Table 1**. In the MDD group (n=206), 99 participants had high depression scores (HAM-D > 17); 60 participants scored in the medium range (HAM-D 14-17) and 47 patients scored in low range (HAM-D <14). All participants in the low HAM-D group were currently taking antidepressants. For 89% of participants, we also had SCID-interview data on the presence or absence of a current MDD episode (**Figure S1C**). Our ability to detect the immunophenotypes described here was likely strengthened by our study eligibility criteria, which excluded people outside the age range 25-50 years and those with major medical problems or substance misuse. All procedures were approved by an independent research ethics committee (National Research Ethics Service East of England, Cambridge Central, UK; approval number 15/EE/0092). All participants provided informed consent in writing and received £100 compensation for taking part.

Participants completed the following clinical assessments and self-report questionnaires: Hamilton Depression Rating Scale; Beck Depression Inventory; Chalder Fatigue Scale; Snaith-Hamilton Pleasure Scale; State-Trait Anxiety Inventory; Childhood Trauma Questionnaire; and Life Events Questionnaire. Venous blood samples were taken for measurement of hematology cell counts; lipid profile; flow cytometry; high-sensitivity C-reactive protein and plasma interleukin-6, see **Supplementary Methods**. This analysis included 206 patients with MDD and 77 matched controls. For further details of recruitment, participant exclusion and phenotyping, see **Supplementary Methods**.

Flow cytometry (one sample per participant) was performed on fresh PBMCs using live-dead stain and antibodies against CD3, CD4, CD8, CD19, CD56, CD14 and CD16 (see **Table S1**). Data were manually gated blind to participant group according to the strategy in **Figure S2**. Absolute cell counts from the haematology panel were multiplied by percentage multipliers derived from the flow cytometry data (**Table S2**) in order to generate estimated absolute cell counts for neutrophils, eosinophils, basophils, red blood cells, platelets, CD4^+^ T cells, CD8^+^ T cells, classical monocytes, intermediate monocytes, non-classical monocytes, CD16^hi^ NK cells, CD56^hi^ NK cells, NKT cells and B cells. See **Supplementary Methods**.

### Statistical analysis

All analyses were performed in [R] version 3.5.1 (R Core Team, 2018) as per the available code (see **Data availability**). P-values were corrected for multiple comparisons using Benjamini-Hochberg procedure. Group effects were tested using Wilcoxon-Mann-Whitney or *χ*^2^ tests.

‘Residual cell counts’ refer to the residuals from a stepwise linear model for each absolute cell count, with initial model:

*Count ∼ sex + BMI + age + recent infection (yes-no) + current tobacco use (yes-no)*

We used principal components analysis to capture the greatest amount of variability in the data, uninformed by clinical phenotype. To identify the variability in the data which most relates to the clinical group, we performed partial least square discriminant analysis (PLS-DA) using the NIPALS algorithm where MDD vs. control status was the response variable, with the predictor variables being absolute or residual cell counts. Partial least squares regression (PLS-R) was used to identify the cellular predictors of depression severity within the MDD group, with the response variable a matrix of the HAM-D, BDI, Chalder Fatigue and SHAPS scores. For both PLS-DA and PLS-R, the p-value for the predictive ability of the model (pQ2) was estimated by permuting group labels or clinical score sets (5000 permutations). Significant weights were defined as those with an absolute bootstrapped z-score >3 as previously described (*48*). See also **Supplementary Methods**.

To detect any discrete immunophenotypes present in the cellular cell count data, we used Gaussian finite multivariate mixture modelling via [R] package mclust (*49*). This detects the clusters present in the data by using an initial hierarchical agglomeration clustering based on scaled, centered single value decomposition of the 14 cell counts, following by an expectation maximization (EM) algorithm to detect the components (clusters) present in the data. The optimal model type and the optimal number of clusters is determined using the Bayesian Information Criterion (BIC). To detect clusters which are more stable and not sensitive to particular data points, we performed consensus clustering of 5000 randomly selected 90% subsets of the data using [R] package clue (*50*). For **Figure 4**, the number of clusters was predefined as two; for **Figure 5**, cluster number was data-driven. Cluster features were compared: for continuous features, we used Kruskall-Wallis tests for the overall effect of cluster and BH FDR-corrected Conover non-parametric tests for post-hoc comparisons; for categorical variables, we used *χ*^2^ tests for the overall effect of cluster, and BH FDR-corrected *χ*^2^ tests for post-hoc pairwise comparisons. See **Supplementary Methods**.

## Supplementary Materials

Supplementary Methods

Figure S1: MDD participant details

Figure S2: Flow cytometry gating strategy

Figure S3, related to Figure 1B: Spearman correlations between immunological, demographic and clinical variables (MDD cases)

Figure S4, related to Figures 2 and 3: Principal components analysis (PCA) and Partial Least Squares Discriminant Analysis (PLS-DA) of absolute cell counts

Figure S5, related to Figure 4: Binary clustering of MDD cellular immunophenotypes

Figure S6, related to Figure 4: Sensitivity analysis: clustering of residuals following linear models of each cell count including age, sex, BMI, tobacco use and recent infection

Figure S7, related to Figure 5: Data-driven clustering of MDD cellular immunophenotypes

Table S1: Flow cytometry staining cocktail Table S2: Derivation of absolute cell counts

Table S3, related to Figure 1A: Univariate comparison of immunological features in patients with MDD compared to controls

Table S4, related to Figure 4: Clinical, demographic and serological features of the binary clustering of MDD immunophenotypes

Table S5, related to Figure 5: Clinical, demographic and serological features of the four stratified MDD cellular immunophenotypes

Table S6, related to Figures 4 and 5: Correspondence between forced two-way and data-driven clustering

## Acknowledgments

We would like to acknowledge Dominika Wlazly, I-Shu (Dante) Huang, Anviti Vyas, Naghmeh Nikkheslat, Alison McColl and Alex Hatton for blood sample processing and data collection. We thank all study participants, research teams and laboratory staff, without whom this research would not have been possible.

## Funding

This work was supported by the Wellcome Trust [104025]. M Lynall was supported by a fellowship and grant from Addenbrooke’s Charitable Trust, Cambridge and a fellowship from the Medical Research Council (MR/S006257/1). M. R. Clatworthy is supported by the NIHR Cambridge Biomedical Research Centre (Transplant and Regenerative Medicine), NIHR Blood and Transplant Research Unit, MRC New Investigator Research Grant, MR/N024907/1; Arthritis Research UK Cure Challenge Research Grant, 21777), and an NIHR Research Professorship (RP-2017-08-ST2-002). E. T. Bullmore and C. M. Pariante are supported by an NIHR Senior Investigator award. This work was also supported by the NIHR Cambridge Biomedical Research Centre (Mental Health) and the Cambridge NIHR BRC Cell Phenotyping Hub, as well as the NIHR BRC at the South London and Maudsley NHS Foundation Trust and King’s College London, London.

## Author contributions

Conceptualization, E.B., M.E.L., M.R.C., C.P., J.C., V.M., P.C., N.H., W.C.D, D.J., P.B.; Methodology, M.E.L., L.T., M.R.C., E.B., C.P., J.C., V.M., P.C., N.H., W.C.D, D.J., P.B.; Investigation, L.T., L.P., J.B., E.B., C.P., J.C., V.M., P.C., N.H., W.C.D., D.J., P.B.; Formal Analysis, M.E.L.; Visualization, M.E.L.; Writing – Original Draft, M.E.L.; Writing – Review & Editing, L.T., M.R.C., E.B., P.C., N.H., W.C.D., J.C., D.J.; Project Administration, J.B., L.P., E.B., D.J.; Supervision, E.B. and M.R.C; Funding Acquisition, E.B., M.E.L., M.R.C., C.P., J.C., V.M., P.C., N.H., W.C.D, D.J.

## Competing interests

N.H. consults for GlaxoSmithKline and is in receipt of research funding from J&J. P.B., D.J. and W.D. are employees of Janssen Research & Development, LLC., of Johnson & Johnson, and hold stock in Johnson & Johnson.

## Data and materials availability

Further information and requests for resources and reagents should be directed to Mary-Ellen Lynall. The sharing of data used in this study is restricted by the informed consent process. Our data cannot be made available on public repositories but will be shared with other scientifically accredited research groups on request (Apollo Repository https://doi.org/10.17863/CAM.40364). [R] code to support the analyses and figure generation is available on Github (login for peer review: username lynallpeerreview, password lynallreview2019).

