## Supplementary Material for "Peripheral blood cell immunophenotyping reveals distinct subgroups of inflamed depression"

#### Supplemental items


- Supplementary Methods
- Figure S1: MDD participant details
- Figure S2: Flow cytometry gating strategy
- Figure S3, related to Figure 1B: Spearman correlations between immunological, demographic and clinical variables (MDD participants)
- Figure S4, related to Figures 2 and 3: Principal components analysis (PCA) and Partial Least Squares Discriminant Analysis (PLS-DA) of absolute cell counts
- Figure S5, related to Figure 4: Binary clustering of MDD cellular immunophenotypes
- Figure S6, related to Figure 4: Sensitivity analysis: clustering of residuals following linear models of each cell count including age, sex, BMI, tobacco use and recent infection
- Figure S7, related to Figure 5: Data-driven clustering of MDD cellular immunophenotypes
- Table S1: Flow cytometry staining cocktail
- Table S2: Derivation of absolute cell counts
- Table S3, related to Figure 1A: Univariate comparison of immunological features in patients with MDD compared to controls
- Table S4, related to Figure 4: Clinical, demographic and serological features of the binary clustering of MDD immunophenotypes
- Table S5, related to Figure 5: Clinical, demographic and serological features of the four stratified MDD cellular immunophenotypes
- Table S6, related to Figures 4 and 5: Correspondence between forced two-way and data-driven clustering

#### Supplementary Methods:

**Participant recruitment and exclusion:**
Participants were recruited in two cohorts, with both control and MDD participants recruited from five clinical sites in both cohorts (Cambridge, Brighton, Glasgow, London and Oxford). In the first cohort (n=229 of the participants included in this analysis), MDD patients were selected to fit specific sets of entrance criteria: currently depressed and unmedicated; currently depressed and medicated with poor response to antidepressant treatment; medicated with good response to treatment; and healthy controls (*6*). In the second cohort of participants (n=54 of the participants included in this analysis) participants (and healthy controls) were recruited for further study with the intention of identifying both high-CRP and low-CRP MDD participants. However, for this analysis, we included all quality-controlled data, including from participants who only attended for the baseline eligibility visit (questionnaires and blood sampling) and were not taken forward for brain imaging and other investigations. There has thus been no filtering of participants by CRP level for the data shown here.

In total, 311 participants in the BioDep study (across both cohorts) had useable peripheral blood flow cytometry data. For most participants, the hematology panel was acquired from the same venepuncture sample as the flow cytometry data. We excluded from analysis: (1) participants who did not have a hematology panel acquired within 1 month of the flow cytometry venepuncture (2) participants missing >2 of the 7 hematology panel features. This reduced the number of included participants to 283.

One exclusion criterion for the BioDep study was a lifetime history of any serious medical disorder likely to compromise the interpretation of immunological data. Of the 283 participants included in this cohort, 276 participants completed a self-report medical questionnaire and had a complete medications list ascertained. Participants with any condition requiring treatment with regular oral corticosteroids were excluded. Participants with mild inflammatory disorders were not excluded. One participant reported regular NSAID use. The self-report medical questionnaire, which asked about past or current conditions, indicated that 41 participants possibly had a current inflammatory disorder as follows: 28 reported asthma (mostly not requiring treatment: 7 requiring inhalers, none requiring oral medication), seven reported psoriasis (none requiring any topical or oral treatment), three reported type II diabetes mellitus (one diet-controlled only; none insulin-dependent), two reported coeliac disease and two reported arthritis (not requiring any treatment). Conservatively, these participants were deemed to have ‘minor inflammatory disease’ for the analysis presented here. Thyroid stimulating hormone was measured (normal range 0.4-4.5 mIU/L): based on this, 15 participants had evidence of biochemical hypothyroidism and 3 participants had evidence of biochemical hyperthyroidism. 6 participants were euthymic but treated with thyroxine or levothyroxine. TSH levels were not significantly different between controls and MDD participants (MWU $P$ = 0.8)

Exclusion of the 41 participants with ‘minor inflammatory disease’ did not substantially alter the data structure: it remained true that the only cell type increased in MDD following FDR correction was the CD4+ T cell count (unadjusted $P$ = 0.002), with trends towards increased neutrophils (unadjusted $P$ = 0.04) and intermediate monocytes in

MDD (unadjusted $P$ = 0.02). The structure of PC1 for the cellular data remained a weighted average of all cell counts (**Figure S4B**) and was associated with MDD diagnosis ($P$ = 0.02) and HAM-D symptom severity (Spearman $\rho$ = 0.23, $P$ = 0.0005). In the cluster analyses, there were no significant cluster-related differences in the proportion of participants with minor inflammatory disease (**Figure S5B, S7B**).

**Clinical scales used**

- Hamilton Depression Rating Scale (HAM-D) is a practitioner-administered depression scale which uses a structured interview to assess symptoms of depression experienced over the past week. The first 17 severity items are summed to produce a symptom severity score. Scores can range from 0-54 and empirically-derived cut-off scores are as follows (*24*): no depression (0-7); mild depression (8-16); moderate depression (17-23); severe depression (≥24)
- Beck Depression Inventory (BDI) is a 21-item self-report questionnaire which assesses symptoms of depression experienced in the preceding few days, including affective, cognitive, somatic and vegetative (but not anxiety) symptoms, reflecting the DSM-IV criteria for major depression. The maximum score is 63. In depressed participants, scores of 14-19 correspond to mild depression, scores of 20-28 correspond to moderate depression, and scores >28 correspond to severe depression.
- Chalder Fatigue Scale (CFQ11) is an 11-item self-report questionnaire which assesses symptoms of physical and psychological fatigue within the last month, with scores ranging from 0 to 33 (most fatigued).
- Snaith-Hamilton Pleasure Scale (SHAPS) is a 14-item self-report questionnaire which measures anhedonia (difficulties in experiencing pleasure) over the preceding few days (e.g. ‘I would feel pleasure when I receive praise from other people’). Scores range from 0 to 14 and an abnormal score is defined as 3 or more.
- State-Trait Anxiety Inventory (STAI) is a 40-item self-report questionnaire which assesses anxiety. It has two subscales - a State Anxiety Scale (STAI-S) which measures subjective feelings of anxiety and a Trait Anxiety Scale (STAI-T) which evaluates more stable aspects of ‘anxiety-proneness’. Scores for each subscale vary from a minimum of 20 to a maximum of 80.
- Childhood Trauma Questionnaire (CTQ). This self-report questionnaire asks about 28 possible experiences during childhood and teenage years, asking participants to indicate how frequently they experienced each scenario (e.g. ‘I believe I was physically abused’). Three of the questions are indicators of minimization/denial. The scores for the other 25 items are summed to produce the CTQ score, which ranges from 25 (no traumatic experiences) to 125.
- The Life Events Questionnaire (LEQ) is a self-report questionnaire which asks about 14 events or problems which may have occurred in the preceding 6 months (yes-no answers, summed to produce the LEQ score), and, if such events did occur, ‘How bad was that for you?’ (summed to produce the LEQ rating). For each participant, we created a z-score from their LEQ score and a z-score from their LEQ rating, then took the mean of these two z-scores to produce the participant LEQ z-score used in this analysis.

**Peripheral blood measurements**

- Flow cytometry percentage counts were generated from fresh peripheral blood samples at each of the five clinical centers as follows: blood was collected in $K_2$EDTA tubes (BD cat: 367525) by peripheral venipuncture, then added to Leucosep tubes (Cat # 163290, Greiner Bio-One Ltd) containing Histopaque for density gradient centrifugation. Leucosep tubes were centrifuged at 800*g* for 15 minutes at room temperature without a break. The interphase layer containing peripheral blood mononuclear cells (PBMCs) was decanted and washed twice in DPBS (Life Technologies, Cat #14200067). 2 x $10^5$ cells were stained for flow cytometry in a total staining volume of 50μL for 20 minutes at room temperature using the following fluorophore conjugated antibodies: anti-CD3, anti-CD4, anti-CD8, anti-CD19, anti-CD56, anti-CD14, anti-CD16, live-dead, and Fc-receptor block (antibody details in **Table S1**). Samples were fixed with BD CellFix (Cat # 340181) and flow cytometry data were acquired on either a BD LSR II, BD FACSCanto II or BD LSRFortessa, using BD Comp beads (Cat #552843) as single-color compensation controls. Following acquisition, samples were gated in FlowJo® by a single researcher blinded to group identity. The gating strategy is shown in **Figure S2**.
- High sensitivity C-reactive protein (hsCRP) and the absolute hematology counts (red cells, platelets, neutrophils, eosinophils, basophils, lymphocytes, monocytes) were measured centrally by a clinical diagnostics laboratory (Q2 Solutions) for all participants across the 5 clinical centers. For some participants, multiple hsCRP and hematology measurements were collected over the duration of the study – for this analysis, we used the data acquired closest in time to the venepuncture used for flow cytometry analysis.
- For plasma IL-6 measurement, peripheral blood was collected into plasma preparation tubes (BD Cat #362799). Following venepuncture, tubes were centrifuged at 1600*g* for 15 minutes at room temperature. Plasma supernatant was frozen at -80°C. Samples were thawed and IL-6 was measured using the MSD Proinflammatory Panel 1 V-plex kit (K15049D) as per the manufacturer's instructions. All samples were above the lower limit of detection of the assay (0.060 pg/mL). 78 of the 283 participants did not have plasma IL-6 measurements available.

**Calculation of absolute cell counts**

Absolute cell counts for the 14 cell types used in this analysis (see **Figure 1A**) were calculated as follows:

(1) Samples were gated in FlowJo™ according to the gating strategy in **Figure S2**. Gated flow cytometry data were imported to [R] using [R] package flowWorkspace (*51*). The data were used to generate percentage features as outlined in the "multiplier" column of **Table S2**. These "multiplier" counts are proportions which are multiplied with the absolute cell counts from the hematology panel in order to generate absolute cell counts of different cell subtypes. For example, a participant's absolute CD4+ T cell count = proportion of lymphocytes which are CD4+ T cells (the multiplier, derived from flow cytometry) * the absolute lymphocyte count (from the hematology panel).

(2) Missing flow cytometry multiplier counts and hematology cell counts were imputed by multiple imputation under a multivariate normal model using [R]

package mix (*52*) via the function ImputeData from [R] package mclust (*49*). In total, only 0.33% of counts in this analysis required imputation.

(3) Percentage multipliers and absolute counts were tested for batch effects of clinical center (5 centers) using a Kruskal-Wallis (KW) test for each cell count. All the percentage multipliers showed marked batch effects related to clinical center, consistent with the fact that flow cytometry was performed locally at each center. For the hematology panel (local venepuncture but assayed centrally), only the absolute lymphocyte count ($P$ = 0.005) and monocyte count ($P$ = 0.08) showed evidence of a center-related batch effect.

(4) For those features showing significant center effects at $P$ < 0.1 by KW test, the features were Box-Cox transformed to normalize the counts using the [R] caret package (*53*) preProcess function; debatched for the effect of clinical center using [R] limma package (*54*) removeBatchEffect function; then inverse Box-Cox transformed to return the data to their original distribution. To avoid the introduction of bias which may occur with unbalanced groups, no design matrix was included in the removeBatchEffect function.

(5) The debatched multipliers (from flow data) were multiplied by the debatched absolute cell counts (from the hematology panel) as per **Table S2** to generate the 14 absolute cell counts used in this analysis: neutrophils, eosinophils, basophils, red blood cells, platelets, $CD4^+$ T cells, $CD8^+$ T cells, classical monocytes, intermediate monocytes, non-classical monocytes, $CD16^{hi}$ NK cells, $CD56^{hi}$ NK cells, NKT cells and B cells.

These 14 endpoints were selected as the 14 cell subsets which could reliably be discriminated based on the flow cytometry panel and haematology data acquired.

**Further detail of statistical analysis:**

Throughout the paper, 'n' refers to the number of participants, except where it is explicitly stated that it refers to the number of clusters.

**Case-control univariate testing**

The effect of clinical group (MDD v.s. control) on continuous clinical features (**Table 1**) and on median absolute cell counts (**Figure 1A**, **Table S3**) was tested using a Wilcoxon-Mann-Whitney test via [R] coin package (*55*) function wilcox_test. Effect size was calculated as the Z statistic divided by the square root of the sample size. The effect of clinical group on categorical features was tested using $\chi^2$ tests via [R] function chisq.test with simulated p-values. Spearman rank correlations between variables were tested using the [R] package Hmisc rcorr function and displayed using the [R] corrplot package.

**Generation of cell count residuals**

In order to generate 'residual cell counts', absolute cell counts were Box-Cox transformed using [R] caret package preProcess as previously. The [R] MASS package function stepAIC was then used to perform model selection with default parameters. The initial model used was:

*Count ~ sex + BMI + age + recent infection (yes-no) + current tobacco use (yes-no)*

For each of the 14 cell counts, the residuals of the chosen linear model were designated the ‘residual cell counts’ and used directly for further analyses (PCA, PLS and clustering). To allow stepwise model comparison and residual calculation, missing values for BMI (8 data points), tobacco use (4 data points) and the presence of recent infection (6 data points) were imputed at their median or modal value. These imputed values were only used to generate residual cell counts, and not used in statistical comparisons between groups or clusters.

**Principal components analysis**
A principal components approach seeks to capture the greatest amount of variability in the data, uninformed by clinical phenotype. Principal components analysis was performed using [R] function prcomp with zero-centering and unit-variance scaling of data prior to singular value decomposition. Inputs to the PCA were either absolute cell counts or the residuals from linear models of the absolute cell counts.

**Partial least squares analysis**
To identify the variability in the data which most relates to the clinical phenotype, we performed a partial least squares analysis. Partial least squares (PLS) is a statistical technique which finds the linear combinations of ‘predictors’ (here, the 14 cell counts) which most relate to variability in the ‘response’ variable (here, case vs. control status or a matrix of symptom scores). Where a component is found significantly to predict MDD status or its severity, the ‘weights’ in the PLS model indicate which of the different cell types most contribute to this ability of the model to predict the response, allowing interpretation of the model.

Partial least square discriminant analysis (PLS-DA) was performed taking MDD v.s. control status as the response variable, with the predictor variables being absolute cell counts or the residual absolute cell counts following linear modelling as above. PLS-DA was performed using the NIPALS algorithm via [R] package ropls (*56*). We used weighted centering of predictors prior to PLS in order to avoid biases due to unequal group sizes (*57*). The number of significant PLS components was determined using the default ropls method. A component is only added to the PLS model if the P2Y metric (proportion of response explained) is > 1% and Q2Y > 0, i.e. if the predicted residual sum of squares of the model including the new component (estimated by 7-fold cross-validation) is less than the residual sum of squares of the model with the previous components only.

Partial least squares regression (PLS-R) was used to identify the cellular predictors of depression severity within the MDD group. The response variable was a matrix of the following clinical symptom scores: HAM-D, BDI, Chalder Fatigue and SHAPS score. Only MDD participants with complete symptom scores across these scales were included in the PLS-R analysis (n=199). The predictor variables were absolute cell counts or the residual absolute cell counts (described above). We used the default mean-centering and unit-scaling of predictor and response variables and the NIPALS algorithm for PLS as implemented in [R] package ropls.

The cumulative Q2Y reflects the predictive ability of the PLS model. For both PLS-DA and PLS-R, the p-value for the predictive ability of the model (pQ2) was estimated by permuting group labels or clinical score sets (5000 permutations): pQ2 is the proportion of permuted cumulative Q2Y values above the original cumulative Q2Y for the model. We used the following method to test whether particular cell counts contributed significantly to the PLS model (i.e. testing the significance of predictor weights) (1) use [R] package boot to resample the participant data and calculate the PLS model (and associated weights) using the resampled data (1000 resamples). (2) Generate a z-score for the cell count weight by dividing the cell count weight obtained in the original PLS model by the standard error of the bootstrapped PLS weights. (3) Significant weights are defined as those with an absolute z-score >3 as previously described (*48*). PLS plots show only those cell count weights found to be significant by this metric. This method is more conservative than using VIP (Variable Importance in Projection) >1 to select significant weights. All cell counts designated significant in this analysis had VIP>1.

**Further details of mixture modelling and consensus clustering**

To generate consensus partitions of participants into clusters, we first performed Gaussian mixture modeling on the total dataset to select the category of models to be considered (e.g. VVI, EEI), then performed mixture modeling on 5000 randomly selected 90% subsets of the data (selection without replacement) using that model type to generate 5000 data partitions. For each clustering partition generated, we used [R] package clue cl_predict function to predict the cluster membership for the left-out 10% of the data. Finally, we used the cl_ensemble function to find the consensus clustering which minimized the Euclidean dissimilarity of the 5000 clustering partitions generated.

For **Figure 4**, the number of clusters was predefined as two. Mixture modeling of absolute counts with forced binary clustering selected a "VVI" model (diagonal, varying volume and shape). Mixture modeling of the residual counts (which are normalized as part of the linear modeling) selected an "EEI" model (diagonal, equal volume and shape). For data-driven clustering (**Figure 5**), a Gaussian mixture modelling of absolute cell counts selected a "VVI" model with 4 components. We also used an alternative goodness of fit metric - the Integrated Completed Likelihood (ICL) - to test for the optimal model type and component number (*58*). For our data, this selected the same model types and cluster numbers as the BIC metric. For data-driven clustering, the number of components was allowed to vary from 1 to 9 at every iteration of the mixture model and the best model was selected by BIC. Notably, for the data-driven clustering, taking the data set as a whole compared to subsampling and consensus clustering generated the same number of clusters (four).

**Statistical comparison of cluster features**

We compared the clinical and demographic correlates of the four clusters (immunophenotypes) detected. For continuous variables (e.g. cell counts, age, BMI, symptoms scores), we used a Kruskal-Wallis test for the overall effect of cluster on the variable. For variables where the Kruskal-Wallis p-value was <0.05, we used [R] package PMCMRplus to perform post-hoc Conover non-parametric tests for significant differences between every pair of clusters, with Benjamini-Hochberg FDR adjustment for multiple comparisons across all the cluster pairs. The p-values shown on graphs comparing features cluster-by-cluster represent corrected p-values, and non-significant comparisons are not shown. For categorical variables (e.g. tobacco use, current antidepressant use), we tested for the overall effect of cluster on the variable using a $\chi^2$ test with simulated p-values via [R] function chisq.test. Where $P < 0.05$ for the overall $\chi^2$ test, we performed post-hoc testing of which pairs of populations differ. A $\chi^2$ test with simulated p-values was performed for each pair of clusters, and the raw p-values were adjusted across all pairwise comparisons using Benjamini-Hochberg FDR correction. Only pairs with adjusted p-values <0.05 were deemed significant. Where there were only two clusters (**Figure 4**) statistical tests performed to compare clusters were simple $\chi^2$ tests with simulated p-values (for categorical variables) or Mann Whitney U tests (for continuous variables).

**Electronic data management:**
Study data were collected and managed using REDCap electronic data capture tools hosted at the University of Cambridge. REDCap (Research Electronic Data Capture) (*59*) is a secure, web-based application designed to support data capture for research studies, providing: 1) an intuitive interface for validated data entry; 2) audit trails for tracking data manipulation and export procedures; 3) automated export procedures for seamless data downloads to common statistical packages; and 4) procedures for importing data from external sources.

### Supplementary Figures:

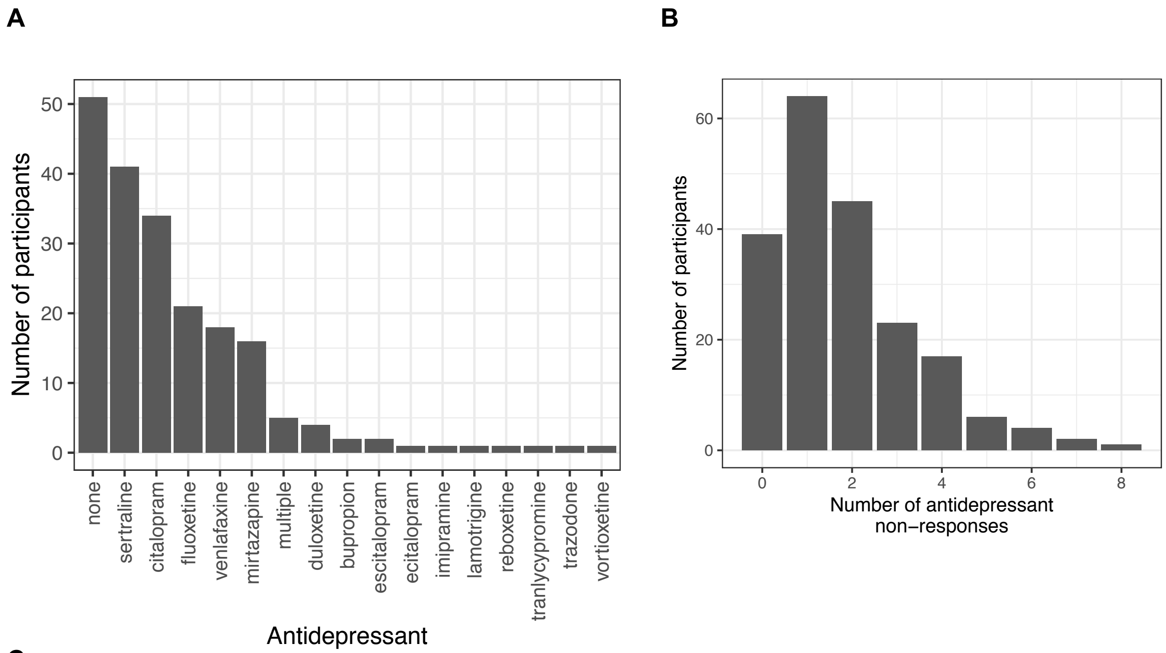


C

| | n | % SCID: MDD current episode depressed | % SCID: MDD current episode HCI depressed | % SCID: not currently depressed | % SCID diagnosis not available | % current antidepressant treatment (n missing data) |
|---|---|---|---|---|---|---|
| HAM-D >17 | 99 | 84 | 1 | 1 | 13 | 56% (4) |
| HAM-D 14-17 | 60 | 46 | 4 | 2 | 8 | 85% (1) |
| HAM-D <14 | 47 | 12 | 2 | 31 | 2 | 100% (0) |

**Figure S1: MDD participant details (A)** Antidepressant medication use at the time of data acquisition for MDD participants. Y-axis indicates the number of participants for each antidepressant shown. ‘Multiple’ indicates use of more than one antidepressant, or antidepressant with adjunctive antipsychotic treatment. **(B)** Number of previous non-responses to antidepressant treatment (<75% response) for MDD participants. **(C)** SCID interview data for all MDD participants (n=206). HCI = high clinical index

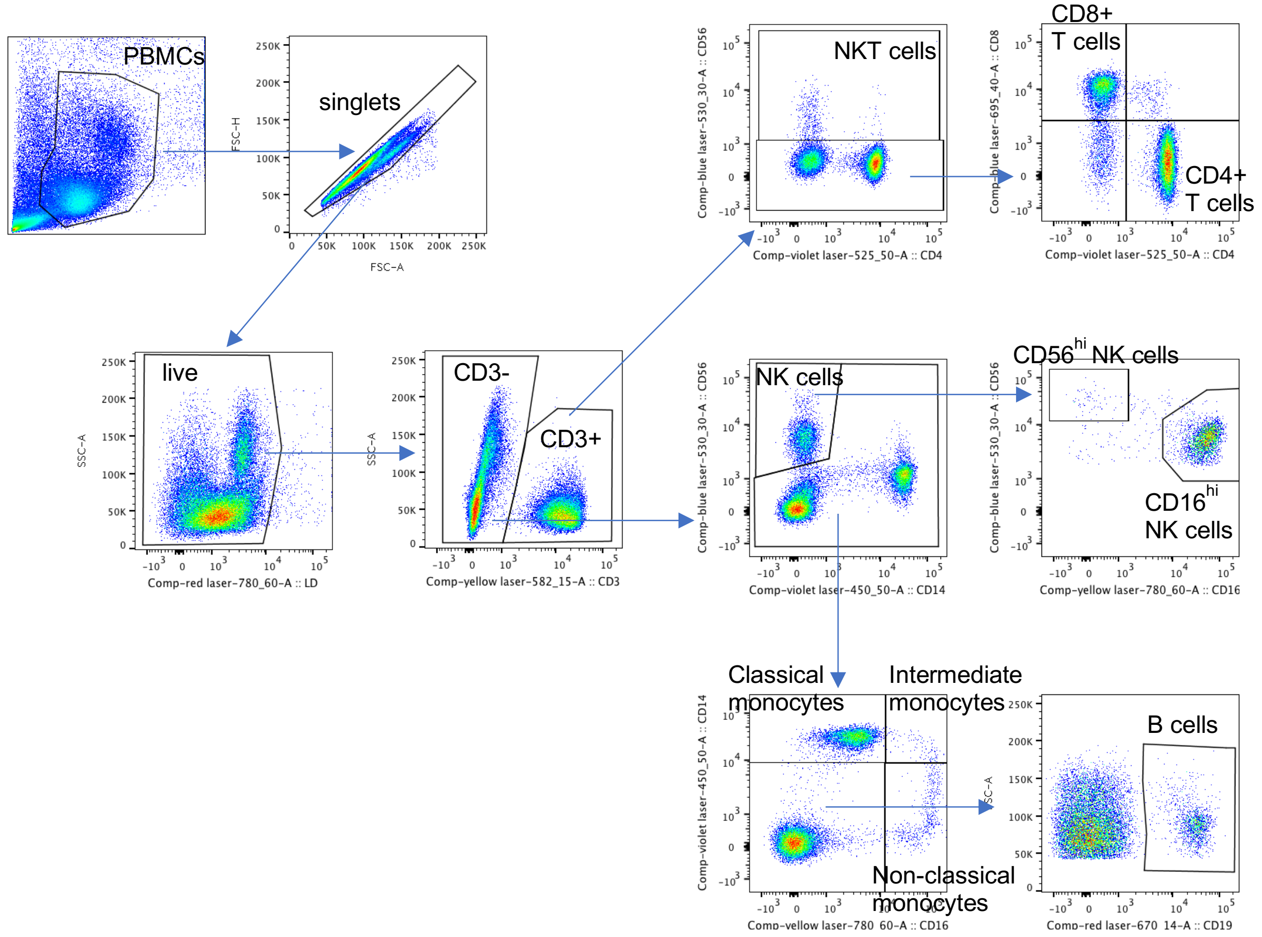


**Figure S2: Flow cytometry gating strategy**

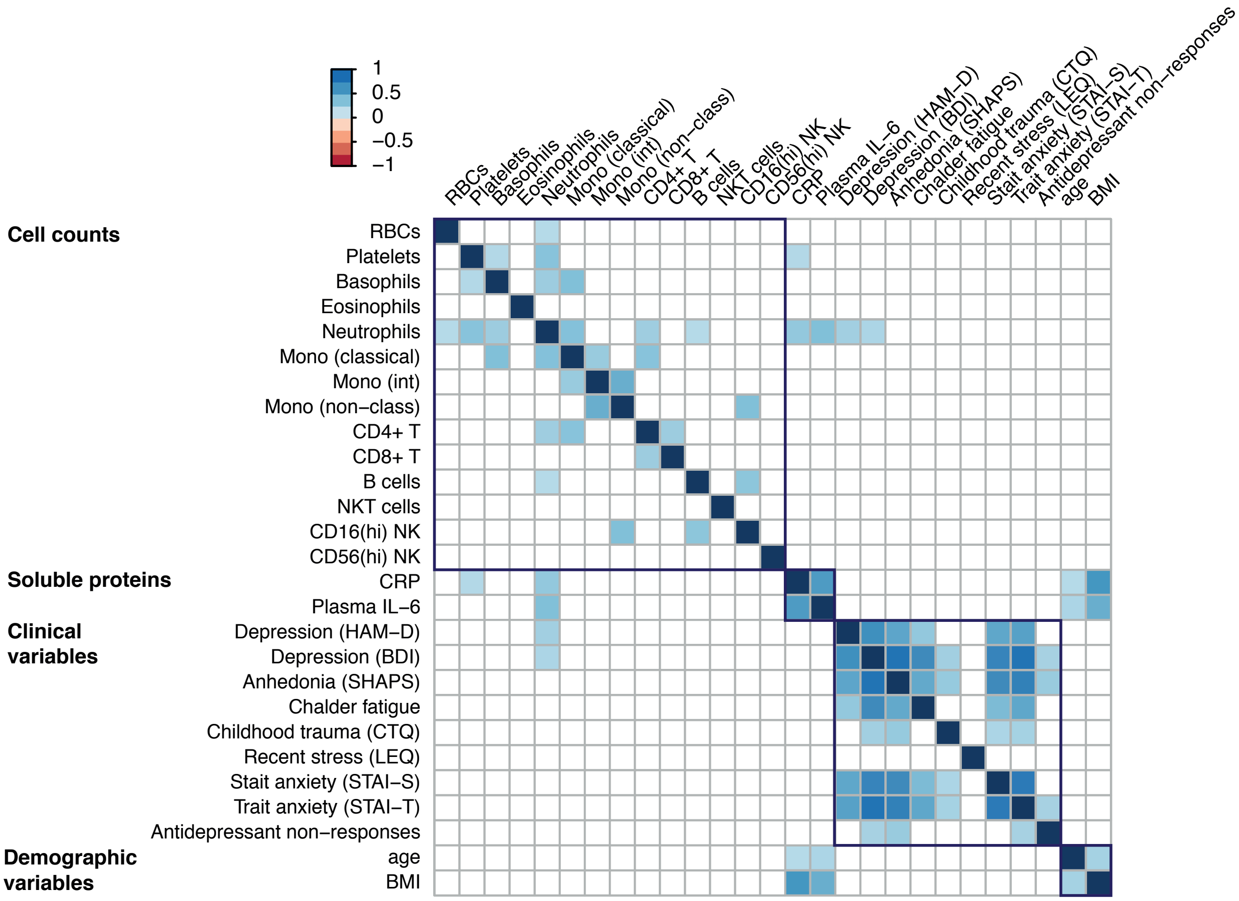


**Figure S3, related to Figure 1B: Spearman correlations between immunological, demographic and clinical variables (MDD participants).**
Data shown for n=206 participants with MDD. Only correlations significant at *P*<0.001 are shown. Correlations including all participants are shown in **Figure 1B**.

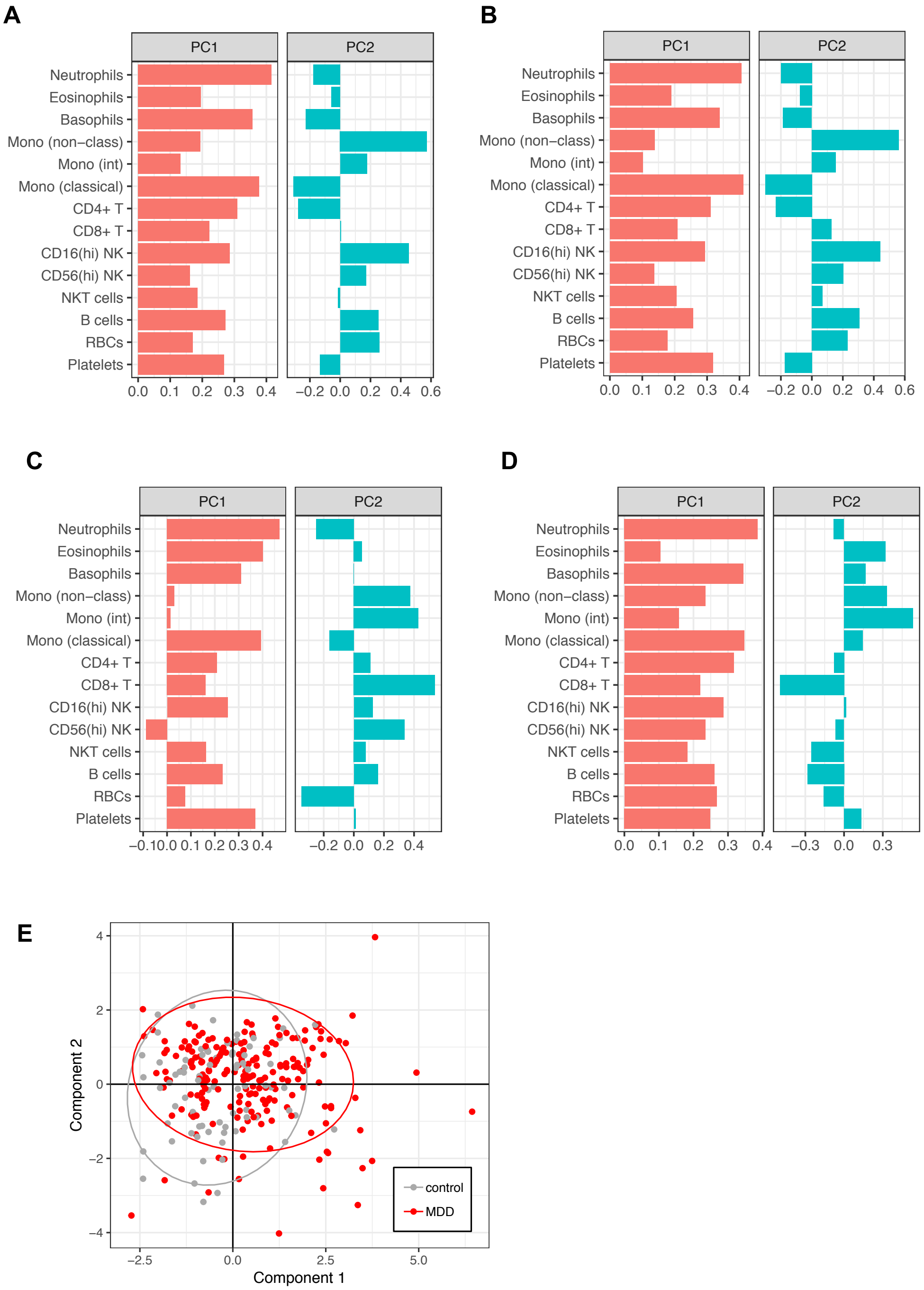


**Figure S4, related to Figures 2 and 3: Principal components analysis (PCA) and Partial Least Squares Discriminant Analysis (PLS-DA) of absolute cell counts. (A)** Absolute counts: cellular PCA for all participants (n=283), see **Figure 2A (B)** Absolute counts: PCA where participants with minor inflammatory conditions have been excluded (n=242) **(C)** Absolute counts: cellular PCA for male participants only (n=88) **(D)** Absolute counts: cellular PCA for female participants only (N=195) **(E)** PLS-DA for major depressive disorder (MDD) vs. controls (n=283). Scatter plot shows the scores on each PLS component for individual participants, where Component 1 is significant. Corresponds to PLS-DA shown in **Figure 3A**.

**Figure S5, related to Figure 4: Binary clustering of MDD cellular immunophenotypes**
(see next page for legend)

**A** Principal components analysis of cell counts (MDD participants only)

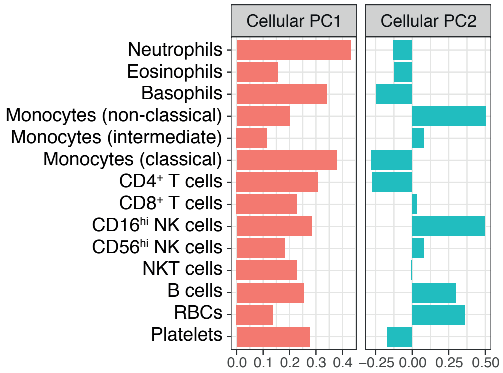


**B** Per-cluster clinical and demographic variables

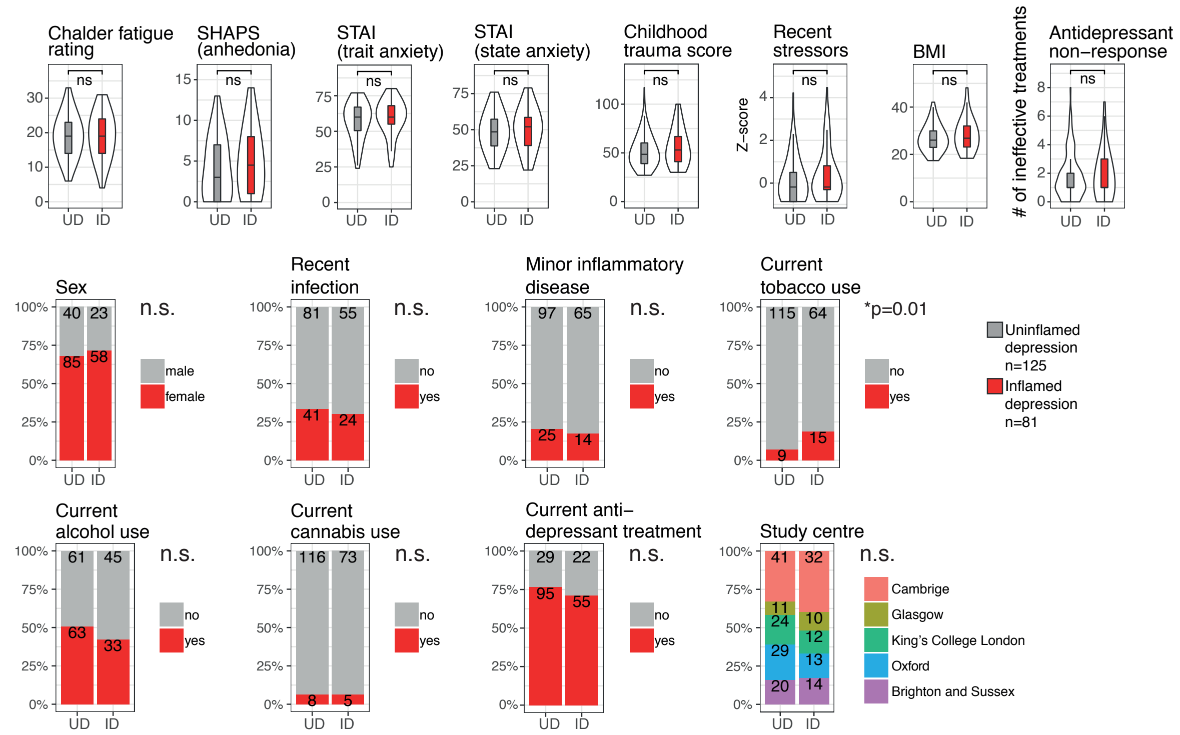


**C** Per-cluster item level scores corresponding to MDD 'typicality' vs. 'atypicality'

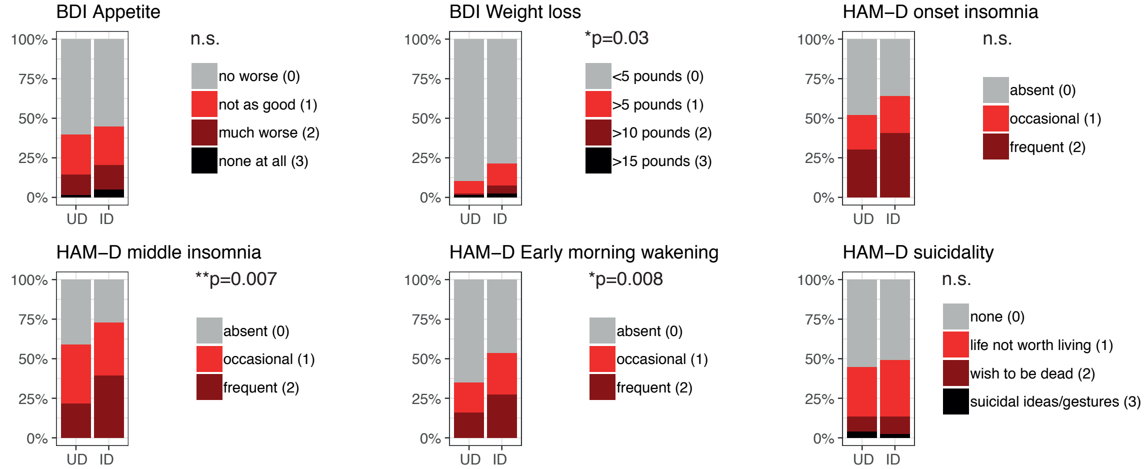

**Figure S5, related to Figure 4: Binary clustering of MDD cellular immunophenotypes**
**(A)** Principal components analysis (PCA) for the 14 absolute cell counts in MDD cases only (n=206). Graphs show the principal component eigenvectors for Principal Component 1 (PC1) and Principal Component 2 (PC2).
**(B)** Forced two-way Gaussian finite mixture modelling of the cellular phenotypes for MDD participants (n=206) identifies two clusters (immunophenotypes) as shown in main **Figure 4**. Boxplots show median and interquartile range for each cluster, with the outer violin shape showing the full distribution of data. Effects of cluster were tested by Mann-Whitney U, **P*<0.05, ***P*<0.01, ****P*<0.01. Bar charts show demographic and clinical variables for each cluster, indicating the percentage of participants in each cluster; bar annotations indicate numbers of participants. The effects of cluster were tested by $\chi^2$. N.s. indicates that the effect of cluster is not significant. See also **Table S4**. SHAPS = Snaith-Hamilton Pleasure Scale, STAI = State-Trait Anxiety Inventory.
**(C)** Participant item-level responses for questions corresponding to MDD 'typicality'. Bar charts indicate the percentage of participants in each cluster giving each response. Effects of cluster were tested by Mann-Whitney U, taking the question responses as an ordinal variable. N.s. indicates not significant.

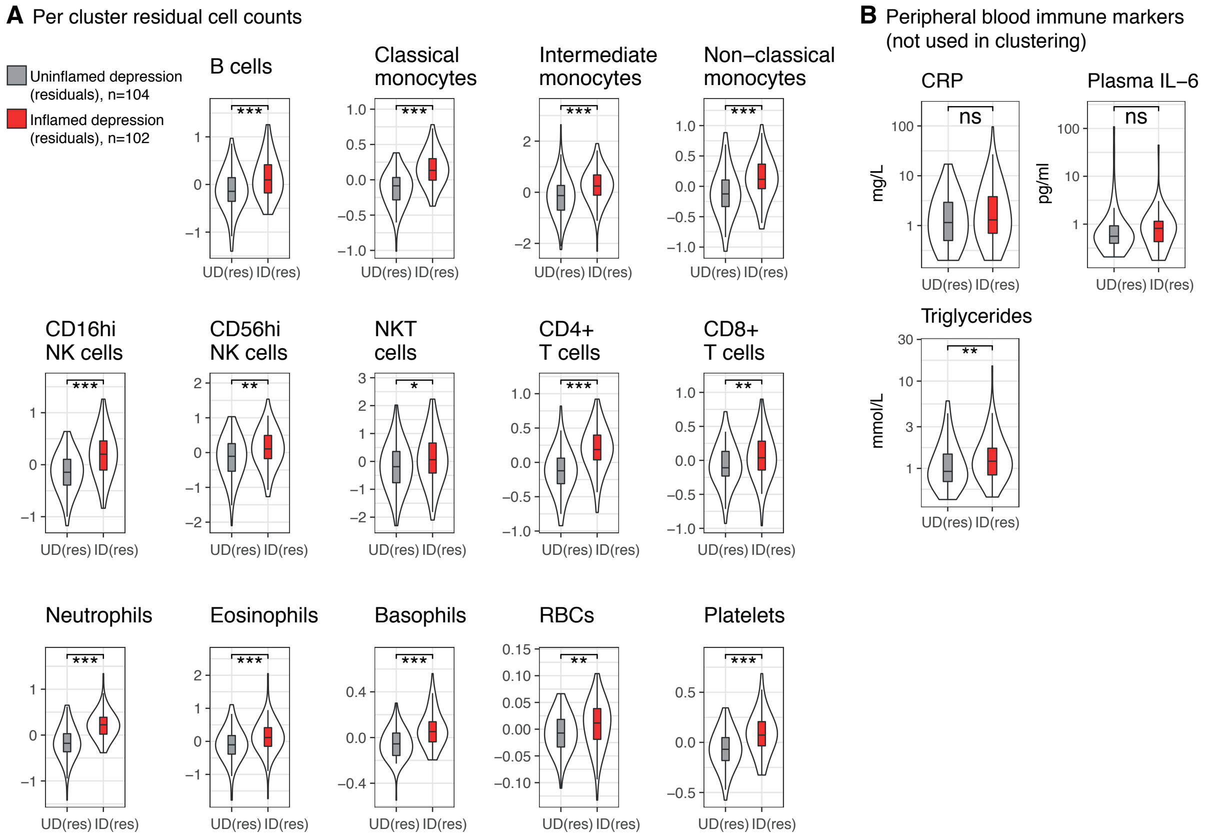


**Figure S6, related to Figure 4: Sensitivity analysis: clustering of residuals following linear models of each cell count including age, sex, body mass index (BMI), tobacco use and recent infection**

Gaussian finite mixture modelling of the residual cell counts from MDD participants once BMI, age, sex, tobacco use and recent infection have been regressed out identified two discrete subgroups (immunophenotypes), uninflamed depression (n=104, grey) and inflamed depression (n=102, red), as shown in main **Figure 4D, 4E**.

**(A)** Boxplots show the median and inter-quartile range of residual cell counts for participants in each cluster, with the violin outline showing the full distribution of data.

**(B)** Peripheral blood markers in each MDD cluster generated from residual cell counts: C-reactive protein (CRP), interleukin-6 (IL-6) and triglycerides. Boxplots show median and interquartile range, with the outer violin shape showing the full distribution of data.

For (A) and (B), clusters were compared by Mann-Whitney U testing: n.s. not significant, **P*<0.05, ***P*<0.01, ****P*<0.001. Data from n=206 participants.

**Figure S7, related to Figure 5: Data-driven clustering of MDD immunophenotypes**
(see next page for legend)

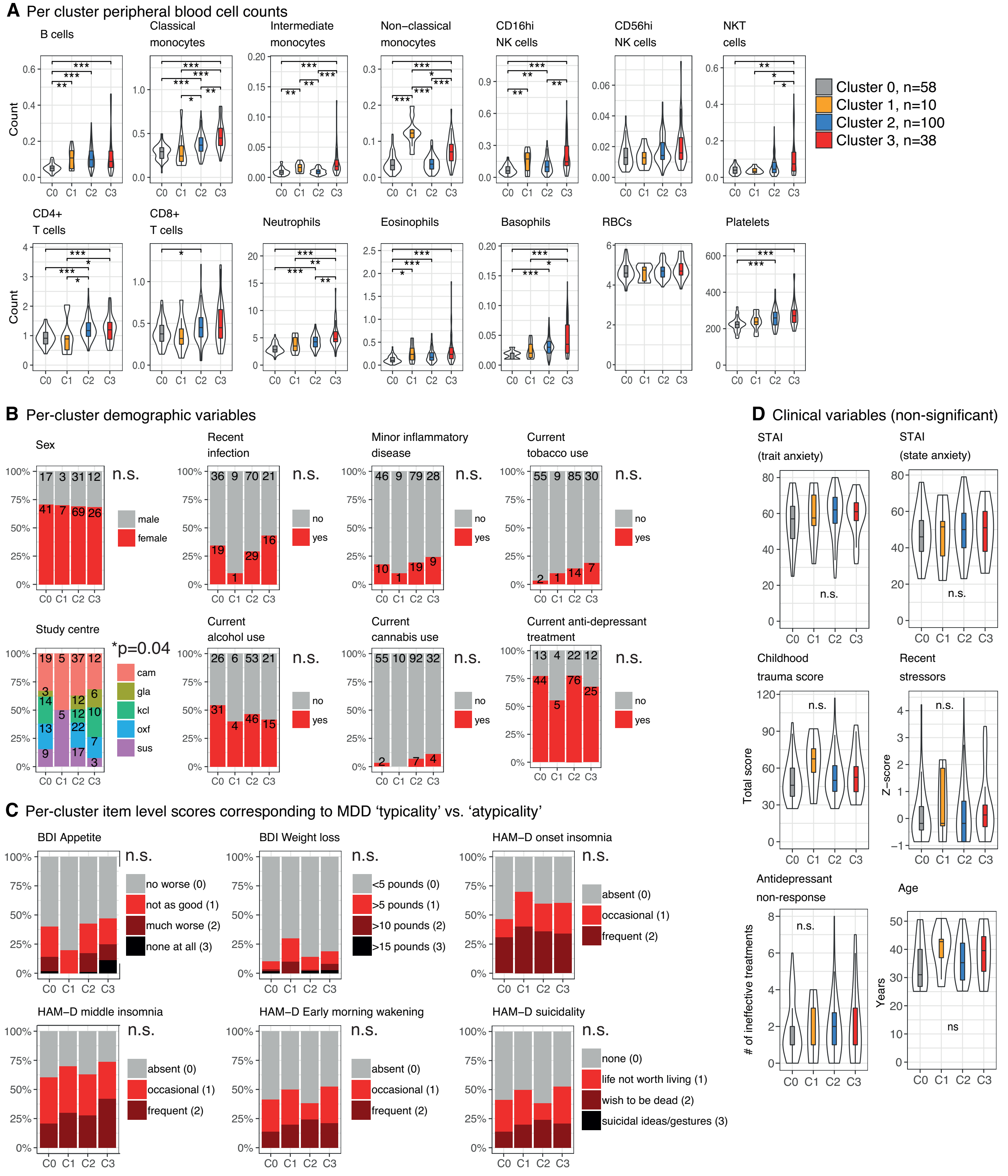

**Figure S7, related to Figure 5: Data-driven clustering of MDD immunophenotypes**
**(A)** Gaussian finite mixture modelling of the cellular phenotypes for MDD participants (n=206) identified four discrete clusters (immunophenotypes) as shown in main **Figure 5**. The arbitrary cluster numbers and colors are used consistently throughout this figure and **Figure 5** to designate each cluster (cluster 0, grey, n=58 participants; cluster 1, orange, n=10 participants; cluster 2, blue, n=100 participants; cluster 3, red, n=38 participants). Boxplots show the median and inter-quartile range of absolute cell counts in each cluster, with the violin outline showing the full distribution of data. The effect of cluster on each feature is tested by Kruskal-Wallis testing. n.s. indicates the overall effect of cluster was not significant at $P<0.05$. Where $P<0.05$ for the overall Kruskal-Wallis test, we performed post-hoc Conover tests for all possible cluster pairs. P-values were corrected for the 6 pairwise cluster-cluster comparisons performed (false discovery rate). Stars indicate adjusted p-values of *<0.05, **<0.01, and ***<0.001. Pairwise comparisons which were non-significant following correction are not shown.
**(B)** Demographic features of each cluster. Bar charts indicate the percentage of participants in each cluster. Annotations on the bars indicate the number of participants in each cluster. N.s indicates that the overall effect of cluster on the feature is not significant by $\chi^2$ testing. * indicates $P<0.05$ by $\chi^2$ testing for the effect of cluster. The results of post-hoc comparisons for each pair of clusters are indicated in **Table S5**.
**(C)** Participant item-level responses for questions corresponding to MDD 'typicality'. Bar charts indicate the percentage of participants in each cluster giving each response. Effects of cluster were tested by Kruskal-Wallis, taking the response as an ordinal variable. As none of the effects of cluster were significant (n.s.) post-hoc comparisons were not performed.
**(D)** Clinical features not differing between clusters (significantly different clinical scores are shown in **Figure 5**). Boxplots show the median and inter-quartile range of the relevant variable for each cluster, violin outline shows the full distribution of data. The effect of cluster on each continuous feature is tested by Kruskal-Wallis testing. STAI = State-Trait Anxiety Inventory.

#### Supplementary Tables:

| Antibody | Company | Catalogue number | Clone | Vol per test (µL) |
|---|---|---|---|---|
| CD14 Pacific blue | BD | 558121 | M5E2 | 1 |
| CD4 V500 | BD | 560768 | RPA-T4 | 1 |
| CD56 AF488 | BD | 557699 | B159 | 1 |
| CD3 PE | BD | 555333 | UCHT1 | 1 |
| CD8 PerCP Cy5.5 | BD | 560662 | RPA-T8 | 1 |
| CD19 APC | BD | 555415 | HIB19 | 1 |
| CD16 PECy7 | BD | 557744 | 3G8 | 1 |
| Zombie NIR | BioLegend | 423106 | | 1 |
| Fc block | Miltenyi | | | 2 |
| DPBS | | | | 40 |

**Table S1: Flow cytometry staining cocktail**

| **Cell type** | **Multiplier (Percentage feature from flow data)** | **Input count** |
|---|---|---|
| Neutrophils | 1 | Absolute neutrophil count |
| Eosinophils | 1 | Absolute eosinophil count |
| Basophils | 1 | Absolute basophil count |
| Red blood cells | 1 | Absolute red blood cell count |
| Platelets | 1 | Absolute platelet count |
| CD4+ T cells | CD4+ T cells / ( CD3+ cells + NK cells + B cells ) | Absolute lymphocyte count |
| CD8+ T cells | CD8+ T cells / ( CD3+ cells + NK cells + B cells ) | Absolute lymphocyte count |
| Classical monocytes | classical monocytes / (classical monocytes + intermediate monocytes + non-classical monocytes) | Absolute monocyte count |
| Intermediate monocytes | intermediate monocytes / (classical monocytes + intermediate monocytes + non-classical monocytes) | Absolute monocyte count |
| Non-classical monocytes | non-classical monocytes / (classical monocytes + intermediate monocytes + non-classical monocytes) | Absolute monocyte count |
| CD16(hi) NK cells | CD16(hi) NK cells / ( CD3+ cells + NK cells + B cells ) | Absolute lymphocyte count |
| CD56(hi) NK cells | CD56(hi) NK cells / ( CD3+ cells + NK cells + B cells ) | Absolute lymphocyte count |
| NKT cells | NKT cells / ( CD3+ cells + NK cells + B cells ) | Absolute lymphocyte count |
| B cells | B cells / ( CD3+ cells + NK cells + B cells ) | Absolute lymphocyte count |

**Table S2: Derivation of absolute cell counts**
Counts for each cell type were obtained by multiplying the value of the “multiplier” column with the “input count”. Where the “multiplier” is non-unity, this is a percentage derived from flow data. The “input count” data are taken from the hematology panel.

| **Humoral factors** | **Control (median, IQR)** | **MDD (median, IQR)** | **P value (MWU)** | **Effect size Z / √ n** |
|---|---|---|---|---|
| High sensitivity CRP (mg/L) | 0.7 (0.4, 1.8) | 1.2 (0.6, 3.3) | *0.003 | 0.18 |
| Plasma IL-6 (pg/ml) | 0.512 (0.346, 0.704) | 0.635 (0.406, 1.071) | 0.04 | 0.14 |
| **Absolute cell counts** | | | | |
| Red blood cells ($x10^6$ /μL) | 4.7 (4.3, 5.0) | 4.7 (4.4, 5.0) | 0.7 | 0.03 |
| Platelets (/μL) | 239 (201, 270) | 242 (217, 284) | 0.7 | 0.08 |
| Basophils ($x10^3$/μL) | 0.020 (0.020, 0.030) | 0.030 (0.020, 0.030) | 0.2 | 0.08 |
| Eosinophils ($x10^3$/μL) | 0.130 (0.080, 0.200) | 0.150 (0.090, 0.250) | 0.09 | 0.10 |
| Neutrophils ($x10^3$/μL) | 3.400 (2.690, 4.180) | 3.875 (2.935, 4.957) | 0.01 | 0.15 |
| Monocytes, classical ($x10^3$/μL) | 0.318 (0.243, 0.405) | 0.348 (0.281, 0.442) | 0.1 | 0.10 |
| Monocytes, intermediate ($x10^3$/μL) | 0.008 (0.004, 0.014) | 0.010 (0.006, 0.015) | 0.02 | 0.14 |
| Monocytes, non-classical ($x10^3$/μL) | 0.037 (0.023, 0.063) | 0.041 (0.024, 0.063) | 0.6 | 0.03 |
| CD4+ T cells ($x10^3$/μL) | 0.934 (0.797, 1.146) | 1.108 (0.862, 1.333) | *0.003 | 0.18 |
| CD8+ T cells ($x10^3$/μL) | 0.430 (0.334, 0.551) | 0.415 (0.320, 0.548) | 0.5 | 0.04 |
| B cells ($x10^3$/μL) | 0.071 (0.042, 0.109) | 0.074 (0.049, 0.121) | 0.3 | 0.07 |
| NKT cells ($x10^3$/μL) | 0.060 (0.030, 0.100) | 0.043 (0.026, 0.075) | 0.08 | 0.11 |
| $CD16^{hi}$ NK cells ($x10^3$/μL) | 0.084 (0.037, 0.136) | 0.099 (0.051, 0.151) | 0.2 | 0.08 |
| $CD56^{hi}$ NK cells ($x10^3$/μL) | 0.015 (0.010, 0.020) | 0.014 (0.010, 0.022) | 0.9 | 0.01 |

**Table S3, related to Figure 1A. Univariate comparison of immunological features in patients with MDD (n=206) compared to controls (n=77)**
P-values for comparison of control vs. MDD by Mann-Whitney U (MWU) testing. * = FDR adjusted $P<0.05$ following correction for 16 multiple comparisons by Benjamini-Hochberg control of false discovery rate. MDD, Major Depressive Disorder; IQR, Interquartile Range; IL-6, interleukin-6. NB. Platelets are included as a ‘cell count’, although not strictly a cell type.

| | **Uninflamed depression (grey)** | **Inflamed depression (red)** | **P-value** |
|---|---|---|---|
| N per cluster (all MDD) | 125 (61%) | 81 (39%) | |
| Cell counts | All cell counts low | Increases in all granulocytes, all monocyte subsets, T cells, B cells and CD16$^{hi}$ NK cells | See **Figure 4C** |
| **Percentage** | | | **P-value ($\chi^2$)** |
| Female | 68% | 72% | 0.7 |
| Current antidepressant use | 77% | 71% | 0.5 |
| Current tobacco use | 7% | 19% | *0.01 |
| Current alcohol use | 51% | 42% | 0.3 |
| Current cannabis use | 6% | 6% | 1 |
| Unemployed | 17% | 33% | **0.008 |
| Recent infection | 34% | 30% | 0.7 |
| Minor inflammatory disease | 20% | 17% | 0.7 |
| Study center | - | - | 0.6 |
| **Median** | | | **P-value (KW)** |
| HAM-D | 16 | 19 | **0.0002 |
| BDI | 22 | 26 | *0.01 |
| Chalder fatigue | 19 | 19 | 0.7 |
| SHAPS (anhedonia) | 3 | 4.5 | 0.1 |
| STAI – state anxiety | 48.5 | 52 | 0.4 |
| STAI – trait anxiety | 60 | 60 | 0.2 |
| Childhood Trauma Questionnaire | 48.5 | 53 | 0.2 |
| Life Events Questionnaire z-score | -0.18 | -0.18 | 0.1 |
| Number of ineffective antidepressants | 1 | 1 | 0.9 |
| Age | 33.6 | 38.4 | *0.01 |
| CRP | 1 | 1.5 | *0.03 |
| Plasma IL-6 | 0.56 | 0.88 | *0.02 |
| BMI | 26.0 | 26.8 | 0.2 |
| Triglycerides | 0.95 | 1.24 | **0.002 |

**Table S4, related to Figure 4: Clinical, demographic and serological features of the binary clustering of MDD immunophenotypes**
MDD, major depressive disorder; BMI, body mass index; IL-6, interleukin-6; BDI, Beck Depression Inventory; HAM-D, Hamilton Depression Scale; SHAPS, Snaith-Hamilton Pleasure Scale; STAI, State-Trait Anxiety Inventory. P values shown are for Kruskal-Wallis (KW) or $\chi^2$ tests of a significant effect of cluster on the variable tested: *$P<0.05$, **$P<0.01$. The associated plots are shown in **Figure 4**. Data from n=206 participants. See also **Figure S5**.

**Table S5: Clinical, demographic and serological features of the four stratified MDD cellular immunophenotypes** (see caption over)

| | **Cluster 0 (grey)** | **Cluster 1 (orange)** | **Cluster 2 (blue)** | **Cluster 3 (red)** | **P-value** |
|---|---|---|---|---|---|
| N per cluster (all MDD) | 58 (28%) | 10 (5%) | 100 (49%) | 38 (18%) | |
| Cell counts | All cell counts low | Increases in non-classical monocytes, CD16$^{hi}$ NK cells and B cells | Increases in adaptive immune cells (B cells and T cells), and some innate immune cells | Myeloid-dominant immune profile. Increase in innate and adaptive immune cells, with particularly marked increases in all monocyte subsets and in neutrophils | See **Figure S7A** |
| **Percentage** | | | | | **P-value ($\chi^2$)** |
| Female | 71% | 70% | 69% | 68% | 1 |
| Current antidepressant use | 77% | 56% | 78% | 68% | 0.3 |
| Current tobacco use | 4% | 10% | 14% | 19% | 0.1 |
| Current alcohol use | 54% | 40% | 46% | 42% | 0.6 |
| Current cannabis use | 4% | 0% | 7% | 11% | 0.4 |
| Unemployed | 11% (b) | 60% (a) | 26% | 24% | **0.004 |
| Recent infection | 35% | 10% | 29% | 43% | 0.2 |
| Minor inflammatory disease | 18% | 10% | 19% | 24% | 0.7 |
| Study center | - | - (d) | - | - (b) | *0.04 |
| **Median** | | | | | **P-value (KW)** |
| HAM-D | 15 | 18 | 18 | 19.5 | **0.009 |
| BDI | 18 | 21 | 27 | 25 | **0.002 |
| Chalder fatigue | 17 | 15.5 | 21 | 20 | **0.005 |
| SHAPS (anhedonia) | 1 | 3.5 | 5 | 4.5 | **0.003 |
| STAI – state anxiety | 46 | 51.5 | 50 | 51 | 0.6 |
| STAI – trait anxiety | 57 | 57.5 | 62 | 61 | 0.2 |
| Childhood Trauma Questionnaire | 46 | 67.5 | 50 | 52.5 | 0.08 |
| Life Events Questionnaire z-score | -0.18 | -0.18 | -0.18 | 0.13 | 0.4 |
| Number of ineffective antidepressants | 1 | 1 | 2 | 1 | 1 |
| Age | 31.0 | 42.7 | 35.3 | 39.6 | *0.02 |
| CRP | 0.60 | 0.95 | 1.50 | 2.45 | ***0.0003 |
| Plasma IL-6 | 0.53 | 0.54 | 0.69 | 0.89 | **0.002 |
| BMI | 25.4 | 23.7 | 26.6 | 31.2 | **0.002 |
| Triglycerides | 0.85 | 0.98 | 1.17 | 1.34 | **0.006 |

**Table S5, related to Figure 5: Clinical, demographic and serological features of the four stratified MDD cellular immunophenotypes** This corresponds to the clustering shown in **Figure 5**. MDD, major depressive disorder; BMI, body mass index; IL-6, interleukin-6; BDI, Beck Depression Inventory; HAM-D, Hamilton Depression Scale; SHAPS, Snaith-Hamilton Pleasure Scale; STAI, State-Trait Anxiety Inventory. P-values shown are for Kruskal-Wallis (KW) or $\chi^2$ tests of the effect of cluster on the variable tested: *$P<0.05$, **$P<0.01$, and ***$P<0.001$. For continuous variables, the post-hoc comparisons are shown in **Figures 5C, 5D, S7D**. For categorical variables, pairwise cluster comparisons which are significant by post-hoc testing following correction for 6 multiple comparison at $P<0.05$ are shown in the table as follows: (a) different from uninflamed depression cluster, (b) different from cluster 1, (c) different from cluster 2, (d) different from cluster 3. For cell counts and post-hoc tests for cell counts, see **Figure S7A**. Data from n=206 participants.

| | | Forced two-way clustering | |
|---|---|---|---|
| | | **Uninflamed depression** | **Inflamed depression** |
| **Data-driven clustering** | **Cluster 0** | 58 | 0 |
| | **Cluster 1** | 0 | 10 |
| | **Cluster 2** | 67 | 33 |
| | **Cluster 3** | 0 | 38 |

**Table S6, related to Figures 4 and 5: Correspondence between forced two-way and data-driven clustering**
